# Single cell analysis of the dorsal V-SVZ reveals differential quiescence of postnatal pallial and subpallial neural stem cells driven by TGFbeta/BMP-signalling

**DOI:** 10.1101/2022.05.20.492790

**Authors:** Guillaume Marcy, Louis Foucault, Elodie Babina, Emeric Texeraud, Stefan Zweifel, Christophe Heinrich, Hector Hernandez-Vargas, Carlos Parras, Denis Jabaudon, Olivier Raineteau

## Abstract

The ventricular-subventricular zone (V-SVZ) is the largest neurogenic region of the postnatal forebrain, containing neural stem cells (NSCs) that emerge from both the embryonic pallium and subpallium. Despite of this dual origin, glutamatergic neurogenesis declines rapidly after birth, while gabaergic neurogenesis persists throughout life. Here, we performed single-cell RNA-sequencing (scRNA-Seq) of the postnatal dorsal V-SVZ for unravelling the mechanisms leading to pallial lineage germinal activity silencing. We identify cell lineage-specific NSCs primed for the generation of neurons or glial cells, as well as a large population of so far uncharacterized quiescent NSCs (qNSC). Pallial qNSCs enter a state of deep quiescence, characterized by persistent TGFbeta/BMP signalling, reduced transcriptional activity and Hopx expression, whilst in contrast, subpallial qNSCs remain transcriptionally primed for activation. Induction of deep pallial quiescence is paralleled by a rapid blockade of glutamatergic neuron production and differentiation. Finally, manipulation of the TGFbeta/BMP receptor Bmpr1a demonstrate its key role in mediating these effects at early postnatal times. Together, our results highlight a central role of TGFbeta/BMP-signalling in synchronizing quiescence induction and blockade of neuronal differentiation to rapidly silence pallial germinal activity after birth.

## Introduction

During development, radial glial (RG) cells residing in the ventricular (VZ) and subventricular zone (SVZ) are multipotent stem cells, generating ependymal cells, neurons and glial cells. Their production is molecularly regulated in space and time. In mammals, glutamatergic neurons are produced by dorsal (i.e. pallial) RG cells, whilst gabaergic neurons are generated by more ventral (i.e. subpallial) regions.

Whilst the vast majority of forebrain neurons are produced before birth, neurogenesis persists in specific brain regions. The ventricular subventricular zone (V-SVZ) is the largest and most heterogeneous germinal region of the postnatal brain (Fiorelli et al., 2015). The ventral and lateral parts of the V-SVZ both derive from the embryonic subpallium, while its dorsal part derives from the pallium (Young et al., 2007). V-SVZ germinal cells are derived from a subset of RG cells, which transiently enter quiescence late embryonically and then gradually reactivate postnatally (Fuentealba et al., 2015; Furutachi et al., 2015). In particular, these cells (defined as neural stem cells (NSCs) or type B cells) continue producing distinct subtypes of olfactory bulb (OB) interneurons throughout life, the vast majority of which are gabaergic.

Such exclusive gabaergic neurogenesis contrasts with the pallial origin of at least a subpopulation of V-SVZ NSCs. Indeed, a majority of RG cells do not undergo a neurogenic to gliogenic switch (Gao et al., 2014), but instead remain able to generate glutamatergic (Glu) progenitors (Donega et al., 2018). These progenitors express Neurog2 and Eomes/Tbr2 and persist until at least 2 months in the mouse (Azim et al., 2017). Although evidence confirms that at least a subpopulation of these cells contributes to glutamatergic neurogenesis within the cortex at birth (Donega et al., 2018), as well as to olfactory bulb neurogenesis during early postnatal life (Winpenny et al., 2011) until young adulthood (Brill et al., 2009), they are vastly outnumbered by gabaergic progenitors. Thus, uncharacterized mechanisms differentially fine-tune the relative abrupt halt in glutamatergic neurogenesis shortly after birth, contrasting with the continuity of gabaergic neurogenesis throughout life.

Single-cell RNA-sequencing (scRNA-Seq) is a powerful approach to unravel the heterogeneity and dynamics of NSCs at adult stages. For instance, prospective isolation of NSCs has been achieved by flow-activated cell sorting of these cells based on the expression of selected markers (Mizrak et al., 2019). Microdissection of the lateral, ventral and medial V-SVZ followed by drop-seq have also been achieved (Zywitza et al., 2018), providing insights into the transcriptional coding of NSCs competence and dynamics. Here, we aimed at complementing these previous studies by focusing on the early postnatal life, a period of transition between embryonic development and adulthood. Further, we focused on a so-far unexplored region of the V-SVZ, i.e. its dorsal compartment where NSCs of both the pallial and subpallial origin coexist (Fiorelli et al., 2015). This allowed us to unravel the transcriptional hallmarks associated with the rapid closure of glutamatergic neurogenesis, while gabaergic neurogenesis persists. We demonstrate that several mechanisms coordinate the rapid silencing of pallial NSCs. These include their transition into deep quiescence and a blockade of neuronal differentiation, which are both regulated by TGFβ/BMP-signalling through Bmpr1a.

## Results

### Analysis of the postnatal dorsal V-SVZ by large-scale single-cell profiling

To gain an in-depth overview of the cellular and molecular heterogeneity of the dorsal V-SVZ, we performed large-scale single-cell profiling from the microdissected dorsal wall of the V-SVZ at postnatal day 12 (**Fig. 1A**). Following quality control of independent replicates (**Fig. S1A & C**), we obtained the transcriptome of 11,279 cells (out of >15,000 raw cell counts) by scRNA-Seq using 10x Genomics protocol and high coverage (median of about 4,500 genes detected per cell). Successful isolation of the dorsal V-SVZ cells were validated by the absence of cells expressing Nkx2-1, Nkx2-3, as well as by minimal expression of Vax1 (67 cells) (Coré et al., 2020). In sharp contrast, expression of more dorsal markers such as Msx1 (1,486 cells), Gsx2 (1,845 cells) and Emx1 (4,082) was detected in a significant proportion of all cells (11.1%, 16.3% and 36.1%, respectively). Further evidence that our strategy was not contaminated by cells of ventral lineages is by virtue of the 4 genes (Rlbp1, Gm29260, Pax6, Dcc) recently identified to be confined throughout the dorsal lineage were consistently detected, whilst the 5 gold standard genes (Adgrl3, Slit2, Ptprn2, Rbms1, Sntb1) delineating the ventral lineage were consistently absent (Cebrian Silla et al., 2021) (**Fig. S1D**).

**Fig. 1.**
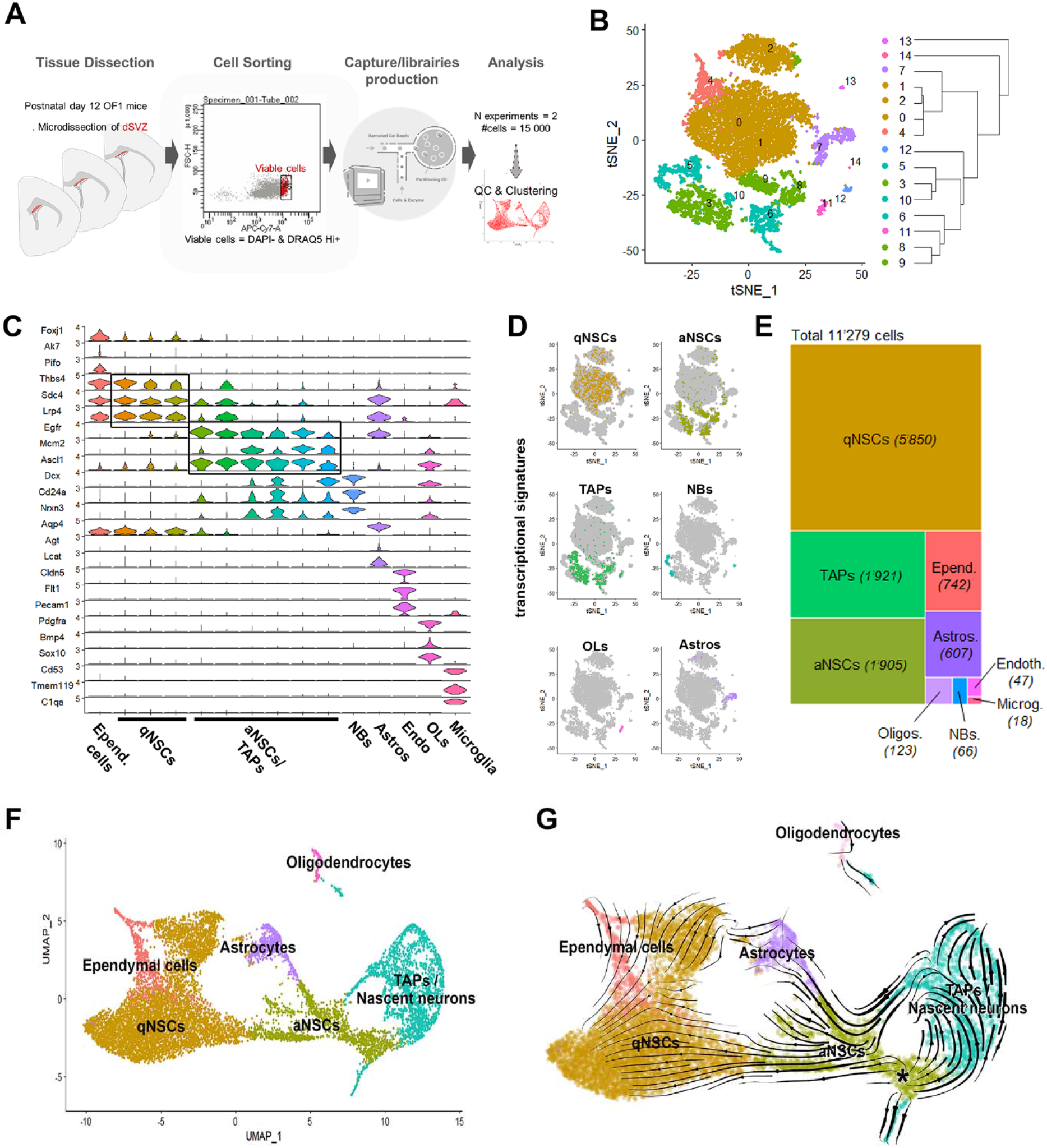
Analysis of the postnatal SVZ dorsal domain cellular composition and differentiation dynamics by large-scale single-cell profiling. **(A**) Scheme of the microdissection and experimental workflow. **(B)** t-SNE projections of major cell types. Color-coding corresponds to cell types shown in D to E. Note that qNSCs represent the larger cell clusters and comprise three clusters of different sizes, which have been grouped in the t-SNE plot (brown). **(C)** Violin plots showing known markers for SVZ cell types. **(D)** Combined expression of key markers on t-SNEs (one combination of markers per major cell type). **(E)** Treemap representing the proportion of the major cell types present in our dataset. **(F)** UMAP with simplified identity annotation. Endothelial cells and microglial cells not shown. **(G)** Pseudotime calculated by RNA velocity highlights the trajectories resulting in the production of qNSCs, glial cells, and TAPs/NBs by aNSCs. The starting point of these trajectories is indicated by an Asterix.

Downstream clustering revealed 15 distinct clusters (resolution 0.5, **Fig. 1B**), amongst which 12 corresponded to cells of the neural lineage. Cell clusters identified were annotated based on the detection of landmark cell type markers, alone (**Fig. 1C**, see also **Fig. S1E-F**) or in combination (**Fig. 1D**). Identification of quiescent NSCs (i.e. qNSCs) was based on the expression of Glast and Prom1 but the exclusion of Egfr and of the ependymal cell marker FoxJ1. Activated NSCs (i.e. aNSCs) were assigned by their expression of Egfr and Ascl1, but exclusion of Dlx1 and Dlx2, whereas transit-amplifying progenitors were defined by an elevated expression of these markers, but low expression of Sp8, Gad1 & Gad2. Finally, we used broad generic markers defining the neuroblast pools (NBs: Dcx, cd24a, and Nrxn3) as well as cells of the oligodendroglial lineage cells (OLglia, Pdgfra & So×10), and concomitant high expression of S100b and Aqp4 for astrocytes (Astros) (**Fig. 1C-D**). These initial analyses identified qNSCs as the prime cohorts of cells captured in our datasets (52% of all cells), followed by aNSCs and TAPs (17% each).

Ependymal cells (6.6%) and Astros (5.4%) were lowly represented. The restricted proportion of oligodendroglia (1%) and NBs (0.5%), underlined the accuracy of our dorsal V-SVZ microdissection. Finally, few non-neural cell types were also present in our datasets (endothelial/mural cells, microglial cells, <0.5%) (**Fig. 1E**).

Altogether, these findings highlight the precision of our microdissection strategy for careful inspection of NSCs at distinct stages of activation, as well as of all their neural progenies, i.e. ependymal, neuronal and glial cells.

### Lineage progression within the postnatal dorsal V-SVZ

We next generated a UMAP with simplified identity annotation for qNSCs, aNSCs and TAPs/NBs (all composed of 3 subclusters) to explore the transcriptional relationship between clusters. Quiescent and activated neural stem cells occupied the center of the UMAP plot, while ependymal cells and neuronal and glial cells appeared as separated clusters in the periphery (**Fig. 1F**, see also **Fig. S2**). In order to gain insights into the major metabolic processes which define the subpopulation of NSCs and their progeny, the top 20 genes associated with previously known markers reflecting state transitions along the neurogenic lineage were overlaid onto the plots (Llorens-Bobadilla et al., 2015). This confirmed the identification of major cell types and revealed parallels with the adult V-SVZ, including: 1) the association of dormancy with elevated glycolytic rates and lipid metabolism in qNSCs (Knobloch et al., 2013); 2) the correlation of ribosomal transcripts with NSC activation (aNSCs); and 3) the rapid sequence of cell cycle initiation, mitosis and neuronal differentiation in TAPs and NBs (**Fig. S3A-B**). Similar Biological Processes (BPs) were highlighted by an over-representation analysis performed on genes up-and down-regulated at transitions between these differentiation stages (**Fig. S3C**). This analysis highlighted the gradual downregulation from qNSCs to TAPs/NBs of genes involved in pluripotency, gliogenesis, as well as glycolytic metabolism, and the inversely correlated upregulation of genes involved in ribosome biogenesis and mitosis (**Fig. S3D**).

We next assessed the cell lineage progression within our datasets by performing non-supervised RNA-velocity lineage trajectory reconstruction (La Manno et al., 2018). In contrast to adult V-SVZ neurogenesis, in which the generation of olfactory neuronal subtypes initiates from qNSCs (Llorens-Bobadilla et al., 2015), postnatal aNSCs were the major pool from which multiple trajectories emerged projecting towards 1) qNSCs, 2) astrocytes and oligodendroglia, and 3) TAPs/NBs (**Fig. 1G**).

Thus, although important similarities can be observed in the NSC differentiation stages and metabolic machineries between postnatal and adult V-SVZ, important differences exist in lineage progression and entry into quiescence between its dorsal and lateral subregions.

### aNSCs are heterogeneous and primed for lineage differentiation

To further understand aNSCs molecular states, we performed a sub clustering analysis. This revealed four different clusters of aNSCs, which were clearly segregated in a UMAP plot (**Fig. 2A-B**), with aNSC3 spatially corresponding to the “starter cell population” identified in the RNA-velocity analysis (see asterisk location in **Fig. 1G**). To determine the main biological differences between aNSC clusters, we performed Gene Ontology analysis on genes enriched in each cluster (aNSC1:522 genes, aNSC2:337 genes, aNSC3:504 genes, aNSC4:525 genes). Genes related to “proliferation” (GO:0006260, GO:0044770, GO:0006281) were amongst those most highly associated to aNSC3, while terms related to “development” (GO:0021537, GO:0030900, GO:0021987), “differentiation” (GO:0010001, GO:0045685, GO:0048709) and “gliogenesis” (GO:0042063) were enriched in aNSC1, aNSC2 and aNSC4 (**Fig. 2C, Data S1**).

**Fig. 2.**
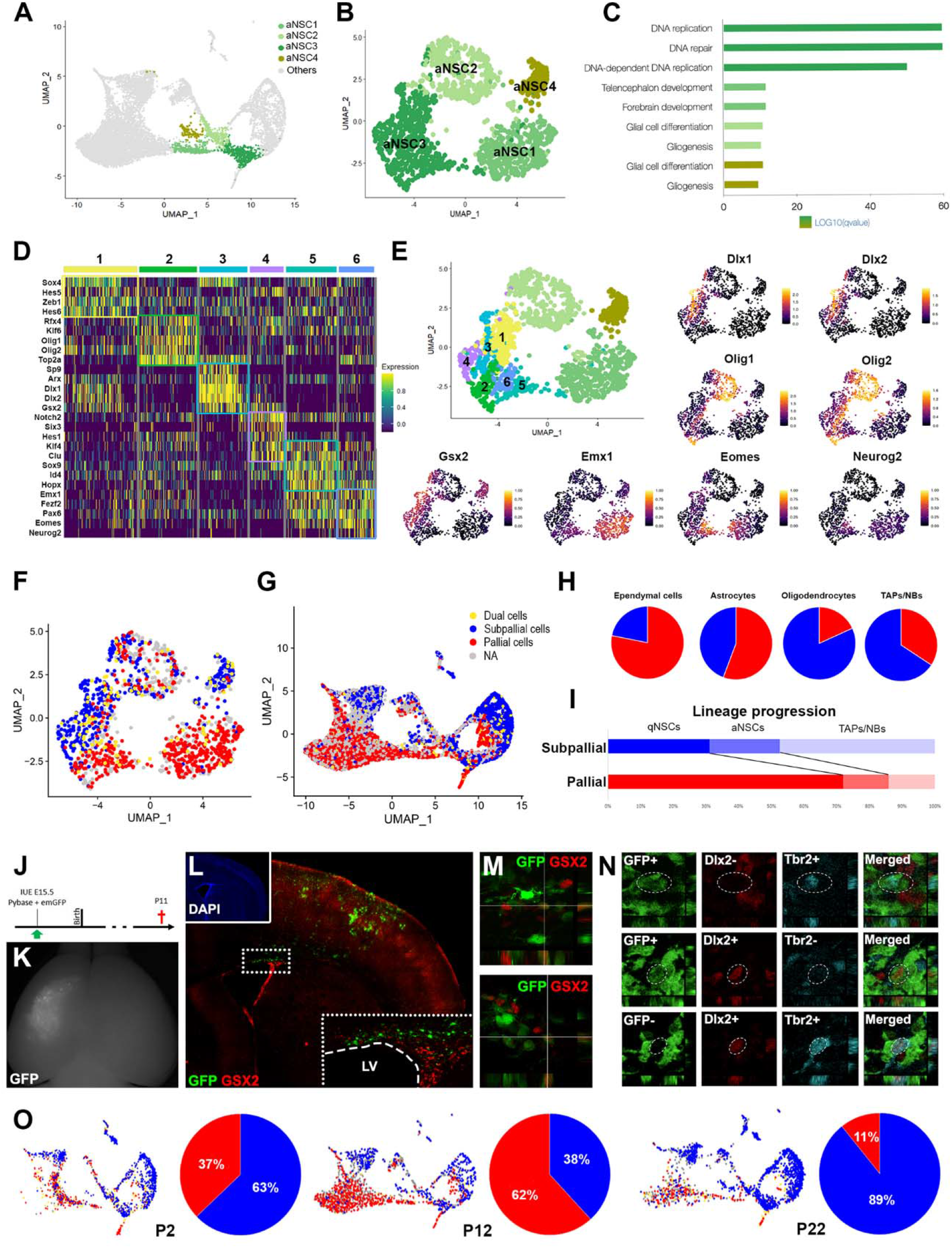
Early priming of pallial and subpallial aNSCs along multiple trajectories. **(A-B)** UMAP plots of complete dataset and aNSCs subsets, with the identity of activated NSC subclusters. **(C)** Bar plot of significantly over-represented Gene Ontology (GO) terms identified by over-representation analysis using genes enriched in all clusters. **(D)** Heatmap showing the top 5 differentially expressed transcription factor/transcriptional regulators for each of the 6 aNSC3 subclusters. **(E)** UMAP plot with identity of aNSC3 subclusters and illustrating feature plots of markers identified in D. **(F-G)** UMAP plot identifying cells expressing pallial (red) or subpallial (blue) transcripts (or both, i.e. dual cells, yellow) within activated NSCs (F) or within the entire dataset (G). **(H)** Percentage of cells expressing pallial (red) or subpallial (blue) transcripts within NSCs progenies. **(I)** Percentage of qNSCs, aNSCs and TAPs/NBs in each lineage. Note the overrepresentation of qNSCs within the pallial lineage. **(J-M)** Fate map of pallial NSCs by electroporation of a transposon GFP plasmid at E15.5, reveals NSCs of pallial origin acquiring subpallial lineage markers at postnatal stages (here Gsx2). **(N)** Postnatal acquisition of subpallial traits by pallial NSCs and their neuronal progeny is further supported by co-expression of Tbr2 and Dlx2 as well as identification of Tbr2+ cells within GAD67GFP+ cells in the dorsal SVZ. **(O)** Analysis of cell proportion at P2, P12 and P22 demonstrate a transient increase of pallial lineage contribution at P12 within the dorsal SVZ, while the subpallial lineage become dominant by P22. See also **Data S1** and **Data S2**.

To gain further insights into this central aNSC3 population (569 cells), we performed additional sub clustering at higher resolution, producing 6 further clusters. Analysis of the differential expression of key transcription factors (TFs) among these clusters (**Fig. 2D & E, Data S2**) supported that the aNSC3 cluster is composed of a mixture of cells primed for differentiation into different lineages, rather than a homogeneous population of multipotent NSCs. In particular clusters 1 and 2 expressed TFs associated with oligodendrogenesis (e.g. Olig1/2), while clusters 4 and 5 expressed TFs associated with neurogenesis (i.e. Dlx2 and Neurog2, respectively). Interestingly, TFs defining subpallial and pallial lineages segregated across clusters 3/4 and 5/6 respectively, demonstrating the coexistence of both lineages within our datasets (**Fig. 2E**).

### Cells of the pallial and subpallial lineages coexist within the dorsal V-SVZ and show different dynamics

Expression of markers for either pallial (Emx1, Neurod1, Neurod6, Neurog2, Tbr1, i.e. 3862 cells) and subpallial lineages (Gsx2, Dlx6, Dlx5, Sp9, Dlx2, Gad1, i.e. 2452 cells), confirmed the coexistence of two spatially segregated lineages within aNSCs (**Fig. 2F**), but also more broadly throughout the entire datasets (**Fig. 2G**). A smaller population of cells expressing both pallial and subpallial markers (“Dual cells”, **Fig. 2F-G**), which were not doublets as revealed by their averaged RNA content (**Fig. S4A**), as well as by “scDblFinder” and “doubletfinder” analyses, suggesting lineage transition for a small number of cells. Quantification of the proportion of cells expressing pallial (Emx1) and subpallial (Gsx2) markers revealed a differential contribution of both lineages to defined cell types (**Fig. 2H**), with with ECs mainly expressing Emx1 pallial marker, astrocytes having equal proportions, whereas OLs and TAPs/NBs mainly expressed the subpallial marker Gsx2 (**Fig. 2I**). Furthermore, comparison of qNSCs, aNSCs, and TAPs/NBs respective proportions between both lineages reveal profound differences. TAPs/NBs were predominant within the Gsx2+ subpallial lineage (i.e. 47% of the cells, compared to 21% for aNSCs and 31% for qNSCs), while qNSCs were predominant within the Emx1+ pallial lineage (i.e. 72% of all cells, with TAPs representing only 14% of all cells) (**Fig. 2H-I**), suggesting different levels of activation/quiescence. The presence of both lineages within the postnatal dorsal SVZ was further confirmed by electroporation of integrative GFP-expressing plasmids in E15.5 pallial NSCs to follow their fate at postnatal times (**Fig. 2J-L**). Immunostaining for Gsx2 at P12 confirmed the emergence of a population of pallial cells expressing subpallial markers (**Fig. 2M**). In addition, colocalization of Dlx2 and/or GAD67::GFP with Tbr2 confirmed that co-expression was not restricted to NSCs, but also includes TAPs/NBs (**Fig. 2N**) in which the proportion of dual cells culminates (i.e. 8% of the cells; **Fig. S4B-C**).

To confirm these results and explore the dynamic of pallial and subpallial lineages within the dorsal SVZ, we produced two additional datasets early after birth (P2) as well as at weaning (P22) (**Fig. S1B**). Results confirmed the presence of cells of the two lineages (**Fig. 2O**). Interestingly, pallial cells peaked at P12, because of a massive increase in the number of pallial qNSCs, but declined by P22 at the expense of cells of the subpallial lineage. Together, these results highlight the coexistence of cells presenting cardinal features of pallial and subpallial lineages within the postnatal dorsal SVZ. Further, our results suggest profound differences in the dynamic of these two lineages, with most pallial cells entering quiescence, while subpallial cells produce a large population of TAPs/NBs.

### Pallial NSCs enter quiescence at postnatal stages within the dorsal V-SVZ and are characterized by Hopx expression

Both the velocity analysis (**Fig. 1G**), see also velocity analysis of pallial and subpallial lineages in **Fig. S4D-F**), as well as the absence of pallial qNSC in our P2 dataset (**Fig. 2O**), suggest their postnatal generation within the dorsal V-SVZ. To confirm this finding, we first integrated our datasets with those of E14.5 and P0 pallial cells (Loo et al., 2019) (**Fig. 3A**). Differentiated mature cells present in both datasets (i.e. microglial cells, endothelial cells and astrocytes) overlapped extensively, validating the robustness of the integration (**Fig. 3B**). Further, significant overlaps were observed for aNSCs/TAPs with progenitor/cycling cell populations, a fraction of TAPs/NBs, and cortical neurons revealing strong transcriptional similarities among these datasets. TAPs/NBs separated in two trajectories corresponding to cortical inhibitory and excitatory neurons, with these cells being rather rare in our datasets due to our microdissection procedure restricted to the dorsal V-SVZ (**Fig. S4G-H**). Remarkably, in sharp contrast to extensive overlaps in all other cell clusters, qNSCs were only observed within our datasets supporting their postnatal emergence (**Fig. 3B**). This was further supported by a ClusterMap analysis highlighting significant similarities in gene expression between all aforementioned cell types, while on the other hand, qNSCs from our datasets were entirely segregated from cell types present within the pallium at E14 or P0 (**Fig. 3C**). To confirm the postnatal generation of qNSCs within the dorsal V-SVZ, we used a label-retaining protocol. At P20, label-retaining progenitors expressing the NSC marker Sox2, but not the ependymal marker S100b, were never observed following BrdU pulse at E14.5 and were rare following a E17.5 pulse. In contrast, they were frequent following BrdU pulses at postnatal times (i.e. P1/3/5), thereby confirming the postnatal appearance of qNSCs (**Fig. 3D**).

**Fig. 3.**
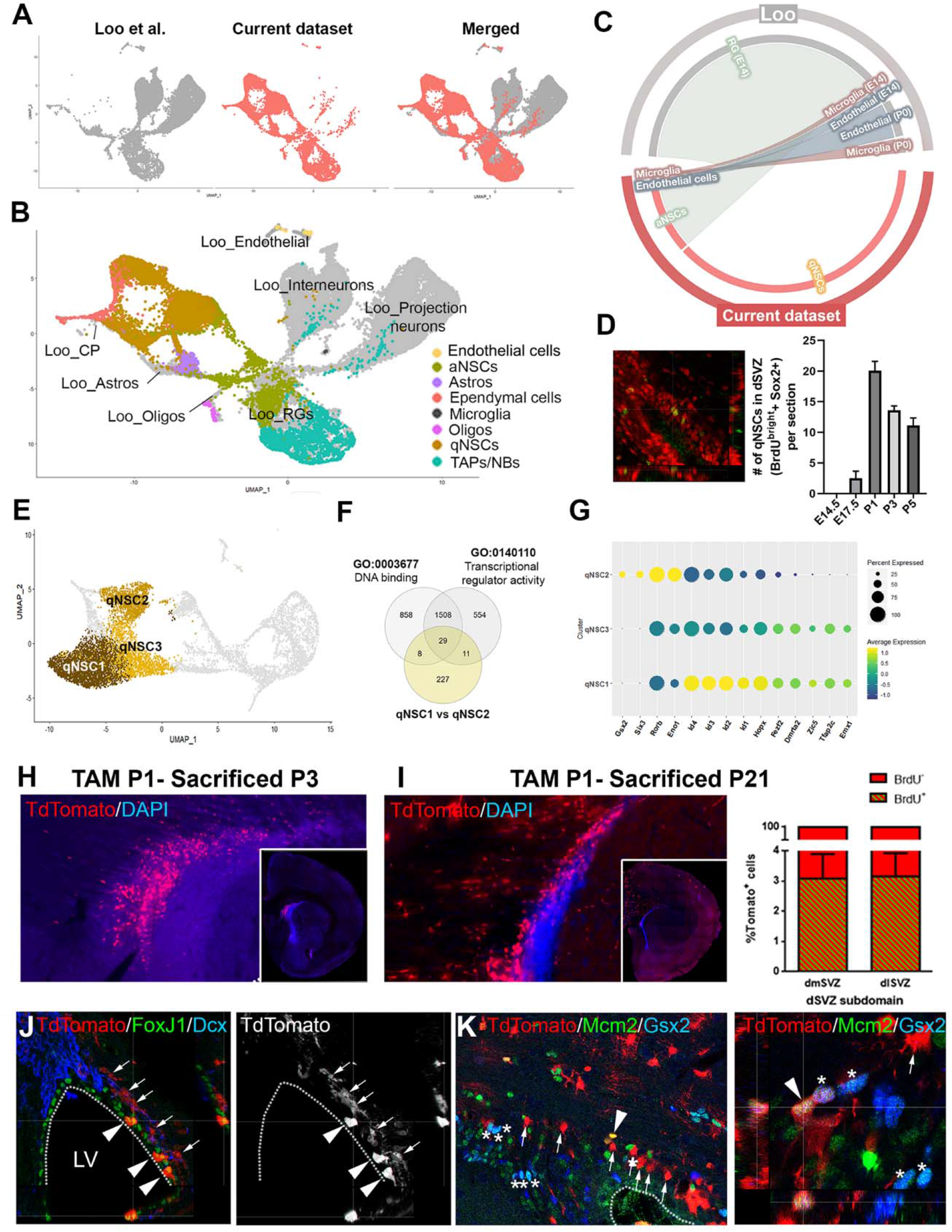
Pallial and subpallial qNSCs are produced postnatally and are distinguished by Hopx expression. **(A)** Integration of current P12 dataset (pink) with previously published dataset of E14.5 and P0 pallium (GSE123335, gray). **(B)** Annotated UMAP depicting the identity of both datasets cell types. Only clusters from the current dataset are colored. See legend for cluster identity. **(C)** Circos plot representing transcriptional correlation between selected cell types of both datasets. Clusters showing transcriptional correlation are relied by line which intensity reflect degree of similarities. **(D)** Quantification of label-retaining cells following BrdU (green) injection at embryonic or postnatal timepoints. qNSCs are defined based on Sox2 expression (red) and S100b exclusion (not shown). **(E)** UMAP highlighting 3 qNSCs subclusters. **(F)** Venn diagram representing DEG between qNSC1 and qNSC2 subclusters and identifying transcription factors/transcriptional regulators (TFs/TRs) identity among them. **(G)** Dot plot analysis revealing that some TFs/TRs are enriched or exclusive to a specific cluster. Among exclusive TFs are the pallial (Emx1) and subpallial (Gsx2) TFs, that define qNSC1 and qNSC2 respectively. Note the expression of several transcriptional regulators including Id proteins and Hopx are enriched in qNSC1. **(H-I)** Fate analysis of Hopx expressing cells. Tamoxifen was injected at P1, then tissue recovered at P3 (H) or P21 (I). Note the large number of recombined tdTom+ cells within the dorsal SVZ at both timepoints. TdTom+ cells have a circular morphology and do not express this marker within the SVZ. BrdU injection 2 hours post Tamoxifen injection only labels 3% of tdTom+ cells at P3 and P21 (arrows), indicating that these cells have entered quiescence and have not resumed cell cycle. **(J)** A small population of tdTom+ cells are ependymal cells and express FoxJ1, while may tdTom+ cells remain away from the ventricular wall and do not express GFAP (not shown), nor the neuroblast marker DCX. (K) tdTom+ and Gsx2+ cells are present within the dorsal SVZ and only minimally overlap. While Gsx2+ and tdTom+/Gsx2+ cells are frequently Mcm2+ (asterisks), tdTom+ cells rarely express this marker further supporting deep quiescence (arrows).

To identify the specificities of pallial and subpallial qNSCs (qNSC1 and qNSC2, respectively, **Fig. 3E and S5A**), we next performed differential gene expression analysis. Focusing on TFs, we confirmed that qNSC1 express higher pallial identity markers such as Emx1, as well as of the transcriptional regulators Hopx, and the Id protein family (**Fig. 3F-G and S5B**). This correlated with an overall reduced transcriptional activity, as reflected by a reduction of 12% in the number of genes detected in qNSC1, when compared to qNSC2 (i.e. 3762 vs. 4276 genes, respectively, **Fig. S5C**). In contrast, qNSC2 showed exclusive expression of subpallial markers including Gsx2, as well as higher expression of Rorb, Eno1 and Six3, which have previously been associated with progenitor proliferation (Appolloni et al., 2008) (**Fig. 3G, Fig. S5B**). The enriched expression of Hopx in pallial qNSCs (qNSC1) is of particular interest given its involvement in integrating TGFβ signalling in other systems (Jain et al., 2015), together with its previously reported expression in the postnatal dorsal V-SVZ (Zweifel et al., 2018) as well as in adult qNSCs (Codega et al., 2014). We took advantage of HopxCreERT2 mice to fate map pallial qNSCs at postnatal stages and confirmed the presence of Sox2(+) qNSCs within the SVZ at P3, that were negative for the ependymal marker FoxJ1 (**Fig. 3J**). Fate mapping 3 weeks after birth, confirmed their persistence (P21, **Fig. 3I**) within the SVZ, although their number decreased and became gradually restricted to the rostral-most region of the V-SVZ. Furthermore, BrdU injection, as well as immunodetection of proliferation markers (Mcm2) confirmed that most fate-mapped cells quickly exited the cell cycle and remained quiescent (**Fig. 3I, K**). Transcriptional analysis at this later timepoint confirmed the presence of pallial qNSCs (qNSC1), as well as of their reduction (-60% from P12 to P22, **Fig. 2O**) while the proportion of subpallial qNSCs remained stable. Finally, BrdU injection, as well as immunodetection of proliferation markers (Mcm2) confirmed that most fate-mapped cells quickly exited the cell cycle and remained quiescent (**Fig. 3J & K**).

### Pallial and subpallial NSCs are defined by distinct levels of quiescence and TGFβ signalling

We next used our transcriptomic datasets to characterize the transcriptional profile of Hopx expressing cells. Identification of differentially expressed genes among Hopxhigh (mostly pallial/qNSC1 cells) and Hopxlow (mostly subpallial/qNSC2 cells, **Fig. S5D**) expressing cells (731 cells Hopxhigh = Hopx>2.5, 651 cells Hopxlow = Hopx<0.4) resulted in the resolving of key ontological categories. Interestingly, Hopxhigh expressed genes involved in “negative regulation of DNA binding transcription factor activity” (GO:0043433), “negative regulation of cell development” (GO:0010721) and “TGFβ signalling pathway” (GO:0007179). In sharp contrast, Hopxlow expressing cells expressed genes involved in “stem cell differentiation” (GO:0048863), “cell maturation” (GO:0048469), and various metabolic pathways, suggesting primed activation (**Fig. 4A, Data S4**). Such primed activation was further supported by a higher number of detected genes when compared to Hopxhigh cells (**Fig. S5E**) as well as by a GSEA analysis, showing that, while no gene sets were associated to Hopxhigh cells, gene sets important for NSCs activation were associated with Hopxlow expressing cells. In particular, they included “mitochondrial biogenesis”, “lipid metabolism”, “fatty acid metabolism”, all defining early stages of NSCs activation (**Fig. 4B, Data S5**). In addition, markers of deep G0 phase (cell cycle inhibitor p27) were enriched in qNSC1 compared to qNSC2, further supporting the state of deep quiescence of those cells (Marqués-Torrejón et al., 2021, **Fig. S5F**). In contrast markers of shallow quiescence (e.g. Cd9) were enriched in qNSC2, paralleled by the enriched expression of markers of activation (Six3, Egfr, **Fig. S5B & G**), as well as of Vcam1, a protein acting as an environmental sensor to regulate V-SVZ lineage progression (Kokovay et al., 2012), and Cpt1a, previously shown to be involved in regulating NSC activity, through the activation of fatty acid metabolism (Knobloch et al., 2017).

**Fig. 4.**
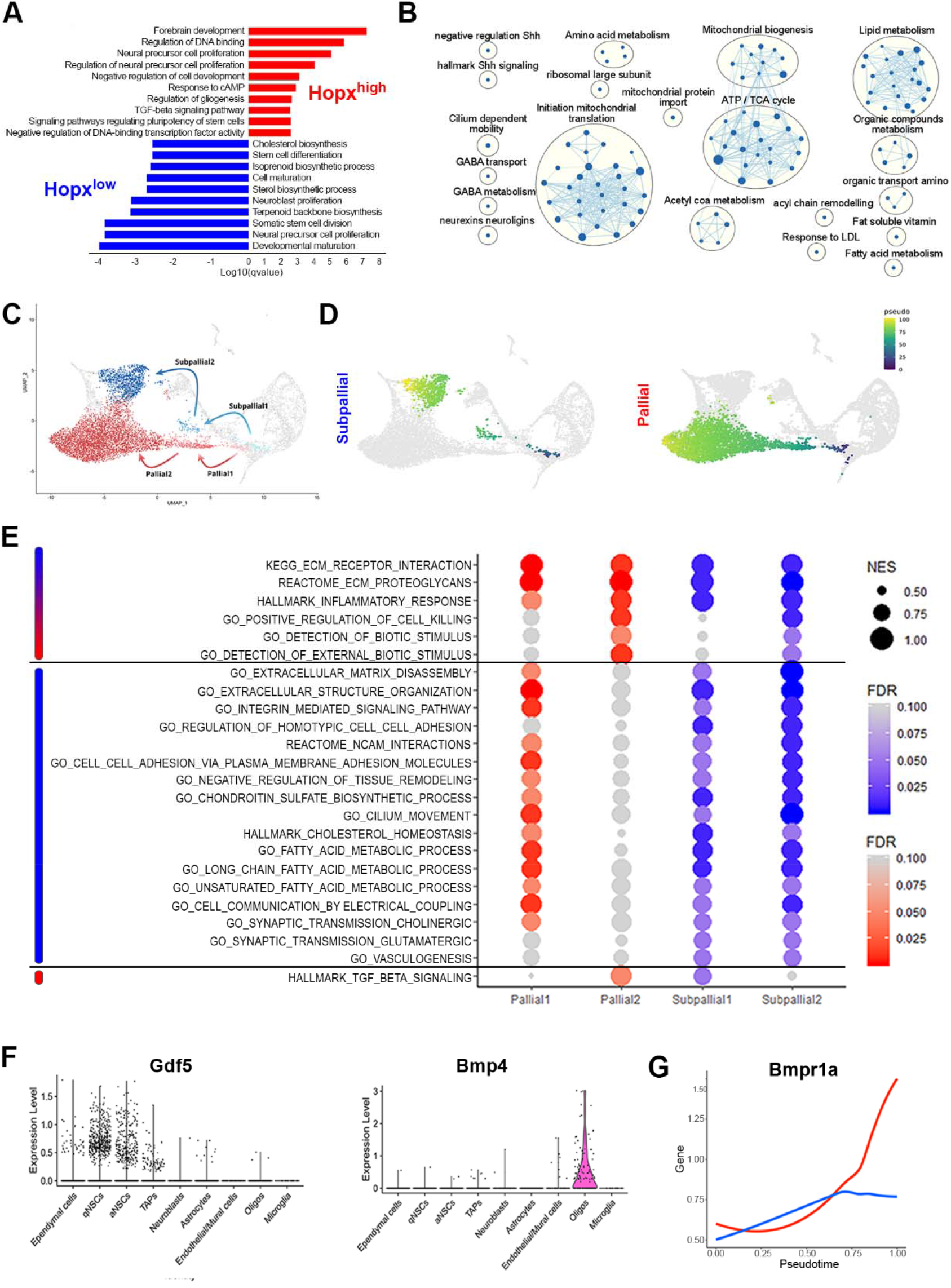
Hopx and TGFβ/BMP signalling define pallial qNSCs quiescence/senescence. **(A)** Bar plot of significantly over-represented Gene Ontology (GO) terms identified by over-representation analysis using genes enriched in Hopx^high^ (red) or Hopx^low^ (blue) cells among qNSCs. **(B)** Hierarchical network of GO terms identified by gene set enrichment analysis using genes enriched in Hopx^low^ (blue) cells among qNSCs. Genes enriched in Hopx^high^ cells did not result in any network. **(C)** Umap illustrating the pallial (red) and subpallial (blue) trajectories and transition stages resulting in the production of qNSCs. **(D)** Pseudotime calculated by RNA velocity highlights the trajectories resulting in the production of qNSCs showing a pallial and subpallial transcriptional signature. **(E)** Identification of key cellular states transitions along both trajectories (see pseudotime for subpallial and pallial trajectories), and selected GO terms associated to the observed transcriptional changes. Associated GO terms can be grouped in 3 categories, i.e. showing similar dynamics (top, blue and red), enriched in the “subpallial” trajectory (middle, blue), enriched in the “pallial” trajectory (bottom, red). **(F)** Violin plot showing expression of Bmpr1a ligands transcripts Gdf5 and Bmp4 in NSCs and Oligodendrocytes (Oligos), respectively. **(G)** Pseudotime analysis of Bmpr1a expression in the pallial and subpallial trajectories. See also **Data S3-5**.

Interestingly, pallial (Hopxhigh) and subpallial (Hopxlow) qNSCs emerged through distinct aNSCs populations. Indeed, aNSC1 and aNSC4 appeared to bridge aNSC3 with pallial and subpallial qNSCs, respectively (**Fig. 4C**, see also **Fig. 2A** and velocity analysis of pallial and subpallial lineages in **Fig. S4F-H, Data S3**). To gain insight on molecular changes defining these two trajectories, we extracted DEGs at each transition stage, and we used GSEA analysis to identify gene sets based on 3 criteria: 1) similar expression within both trajectories, 2) predominance in the subpallial trajectory, 3) predominance in the pallial trajectory (**Fig. 4D &E**). Only a limited number of gene sets showed a similar induction within both trajectories, among which gene ontologies related to ECM receptor interaction (KEGG:M7098), as well as detection of biotic stimuli (GO:0009595, GO:0098581), reflecting the increased anchorage of qNSCs within the niche and interaction with their microenvironment (Codega et al., 2014). Similarly, gene sets related to inflammation (Hallmark:M5932) and regulation of cell killing (GO:0031343) were equally upregulated implying that at least a fraction of qNSCs may undergo cell death.

Genes showing specific upregulation within the subpallial trajectory were more numerous and often associated with biological functions important for NSCs homeostasis. For example, these included multiple genesets related to cell adhesion (i.e. « GO:0034110 regulation of homotypic cell-cell adhesion », « GO:0098742 cell-cell adhesion via plasma-membrane adhesion molecules ») (Codega et al., 2014), cilium biology (GO:0003341 cilium movement) (Tong et al., 2014)), fatty acid metabolism (« GO:0006631 fatty acid metabolic process », « GO:0001676 long-chain fatty acid metabolic process », « GO:0033559 unsaturated fatty acid metabolic process » (Knobloch et al., 2013)), synaptic transmission (« GO:0007271 synaptic transmission, cholinergic », « GO:0035249 synaptic transmission, glutamatergic »), and vasculogenesis ((Lange et al., 2016); GO:0001570 vasculogenesis). In sharp contrast, only a few gene sets were specifically enriched in the pallial trajectory. The most noticeable was the “hallmark of TGFβ signalling” (Hallmark:M5896), which was downregulated in subpallial cells, while constantly increasing with the pallial lineage (**Fig. 4D-E**). Due to the convergence of both analysis in identifying TGFβ signalling in Pallial qNSCs (i.e. see **Fig. 4A & E**), we next explored the pattern of expression of TGFβ ligands and receptors, as well as their dynamic expression by pseudotime. Our analysis detected Bmp4 and Gdf5 as putative TGFβ ligands expressed by OLs and qNSCs themselves (**Fig. 4F**) and identified Bmpr1a as a likely receptor in integrating TGFβ signalling in the pallial lineage (**Fig. 4G**).

Altogether these results highlight the emergence of two qNSCs populations within the postnatal dorsal V-SVZ. Notably, a large cohort expressing pallial genes and persistent TGFβ signalling enters deep quiescence, whereas conversely, a smaller pool expressing subpallial genes remains primed for activation.

### Postnatal induction of deep pallial quiescence is paralleled by a rapid blockade of glutamatergic progenitors/nascent neurons production and differentiation

To investigate the neuronal output of dorsal NSCs, we isolated cells of aNSC3 neuronal trajectory (i.e. aNSC3 subclusters 4 and 5 from **Fig. 2E** and TAPs and nascent neurons). Following removal of Olig1/2 positive cells, we obtained 1029 cells with a majority (66%) expressing Pax6 and Rlbp1, markers of dorsally born NBs (Cebrian Silla et al., 2021), while almost none (3/1029) expressed markers of ventrally born NBs (Runx1t1 and Vax1), confirming the exclusion of NBs from ventral regions in our datasets. New clustering of these cells followed by cluster identification using neural cell subtype markers (see methods for details) resulted in the generation of a UMAP distinguishing two main lineages corresponding to GLU and GABA cells (**Fig. S6**). Both lineages were subdivided into 4 subclusters corresponding to distinct steps of their differentiation, which we named GLU1 to 4 and GABA1 to 4 (**Fig. 5A**). Integration of these cells with those derived from the isolated postnatal OB (Mizrak et al., 2020), resulted in a clear overlap with cycling cells, and two opposing trajectories towards neuronal populations generated postnatally by the dorsal V-SVZ. The first trajectory corresponded to the GLU’s lineage in close proximity with periglomerular (PG) glutamatergic neurons, whose generation persists during postnatal development, albeit at a waned pace (Brill et al., 2009; Winpenny et al., 2011), whereas the second GABA lineage trajectory projected towards granular cells and periglomerular CR+/TH+ interneurons (**Fig. 5B**). Upon the closer inspection of the enrichment of cell cycle stage markers, interestingly, all GABA subclusters corresponded to cycling cells, suggesting sustained amplification, whereas some GLU cells had exited cell cycle (i.e. GLU4), to become postmitotic (**Fig. 5C**).

**Fig. 5.**
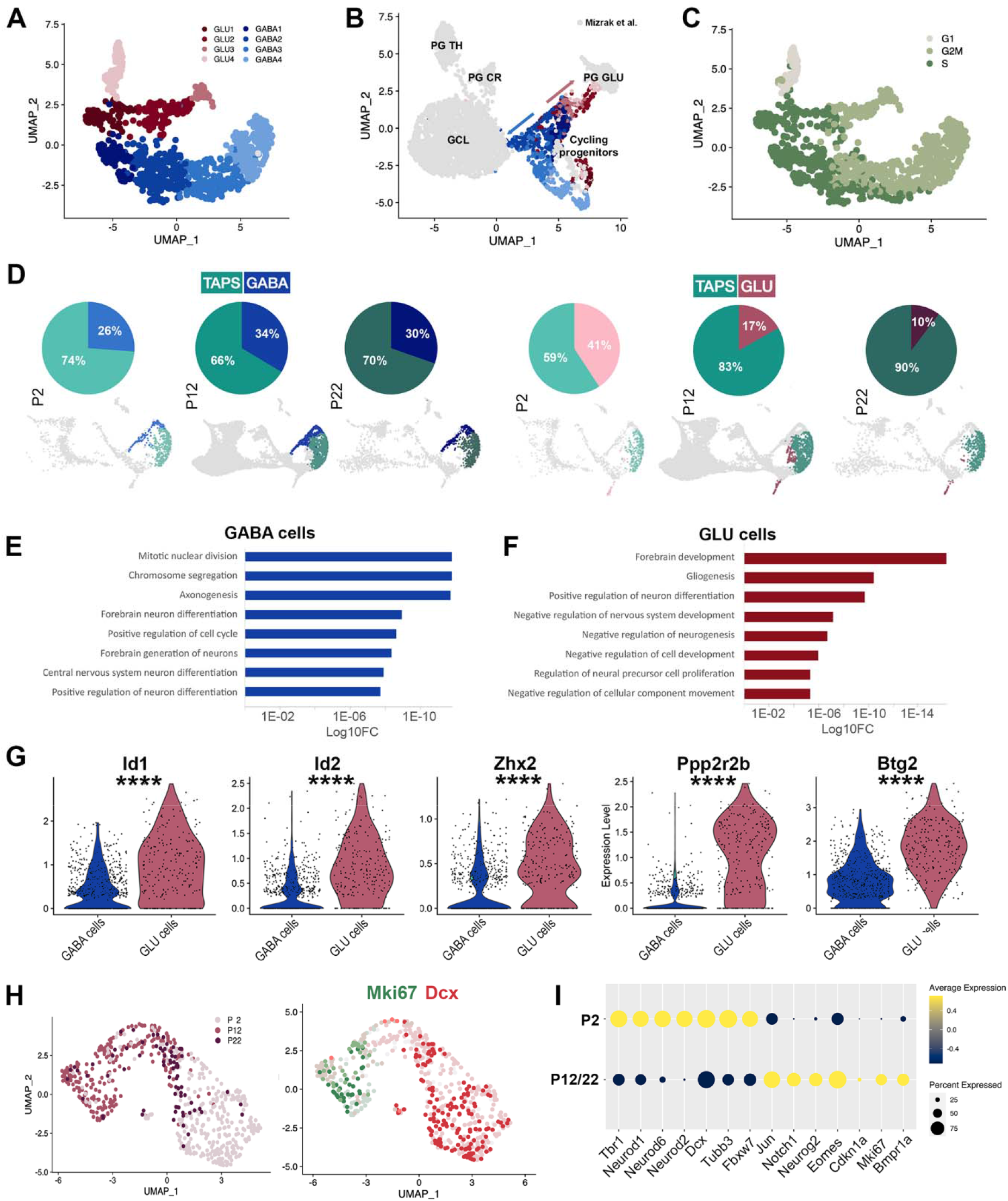
Gradual blockade of pallial lineage neuronal differentiation within the postnatal dorsal SVZ. **(A)** UMAP plot with identity of GLU (deep red gradient) and GABA (deep blue gradient) TAPs/NBs subclusters at P12. **(B)** Integration of current P12 dataset with previously published dataset of postnatally born olfactory bulb neurons (GSE134918, gray). Annotated UMAP depicting the identity of both datasets cell types. Only clusters from the current dataset are colored. See legend in A for cluster identity. Note the presence of two trajectories (arrows) contributing to GABAergic and glutamatergic olfactory bulb neurogenesis. **(C)** Feature plot indicating cycle phase, S.Score and G2M.Score of GLU and GABA cells. **(D)** Pies (and related UMAPs) showing the proportion of GLU cells and GABA cells among TAPs at P2, P12 and P22. Note the rapid decline of GLU cells while GABA cells proportion remain stable. **(E-F)** Bar plots of significantly over-represented Gene Ontology (GO) terms, selected among the top20 categories, identified by over-representation analysis using genes enriched in GABA cells (E) or GLU cells (F). **(G)** Violin plots illustrating selected genes from representative GO terms over-represented in GLU cells. **(H)** UMAP plot of GLU cells at P2, P12 and P22 and Feature plot indicating expression of Mik67 and Dcx. Note that cell cycle exit and neuronal differentiation is observed at P2, but is blocked at later timepoints (P12 & P22). (B) Dotplot illustration enrichment of representative transcripts enriches in GLU cells at early (P2) or late (P12/P22) timepoints. See also **Data S6**.

We next compared the proportion of GABA and GLU cells among TAPs at P2, P12 and P22. While GABA cells remained stable in line with the persistent GABAergic neurogenesis, the proportion of GLU cells rapidly declines (**Fig. 5D**). To investigate the transcriptional correlate of these different dynamics, we identified DEG between cycling GABA and GLU cells. An over-representation analysis revealed key biological processes as differentially regulated between GABA and GLU cells (**Fig. 5E & F**). GO categories associated with mitosis (e.g. “mitotic sister chromatid segregation” (GO:0000070); “chromosome segregation” (GO:0007059); “positive regulation of cell cycle” (GO:0045787), as well as “forebrain neuron differentiation” (GO:0021879) and “regulation of axonogenesis” (GO:0050770), appeared within the top enriched categories in GABA cells (**Fig. 5E** and **Data S6**). In contrast, several GO categories related to “negative regulation of cell development”, “negative regulation of neurogenesis” and “negative regulation of nervous system development” were enriched in GLU cells (**Fig. 5F** and **Data S6**).

These results are in line with the sustained expression of transcriptional repressors (i.e. Id1, Id2, Zhx2;), as well as with the higher expression of anti-proliferation and negative regulator of cells growth genes (i.e. Btg2 and Ppp2r2b, respectively) in GLU cells (**Fig. 5G**).

Finally, we subclustered GLU cells from the P2, P12 and P22 datasets to generate a new Umap. While postmitotic cells were prevalent at P2, their proportion gradually decreased at later timepoints (**Fig. 5H**), illustrating the rapid blockade of GLU cells differentiation. To gain insight into the transcriptional correlates of this blockade, we compared the expression of early and late GLU lineage markers at P2 vs P12/P22 (**Fig. 5I)**. This analysis confirmed enrichment of postmitotic GLU lineage markers at P2 (Tbr1, Neurod1, Neurod2, and Neurod6) in line with the higher expression of immature neuronal markers Tubb3 and Dcx. In contrast, early GLU lineage markers Eomes and Neurog2 were enriched at P12/P22, in agreement with a higher expression of the proliferative markers Cdkn1a and Mki67. Interestingly, Fbxw7 was downregulated at this later timepoint, which together with the enriched expression of Jun and Notch1 may participate in the gradual depletion of GLU cells from the dorsal V-SVZ (Hoeck et al., 2010). Finally, Bmpr1a appeared to be enriched at P12/P22 in a large fraction of GLU cells.

Altogether, our findings indicate that a rapid blockade of glutamatergic neurogenesis parallels the induction of deep quiescence within pallial NSCs, while gabaergic neurogenesis persists in the postnatal dorsal SVZ. Further, persistent high expression of Bmpr1a suggest a role for TGFβ signalling in silencing pallial germinal activity by synchronizing quiescence induction and blockade of neuronal differentiation.

### Manipulation of Bmpr1a signalling modulates dorsal SVZ germinal activity

To investigate the role of Bmpr1a in shaping postnatal dorsal VSZ germinal activity, we performed electroporation of constitutively active, as well as an inactivated form of Bmpr1a (CA Bmpr1a and Δ Bmpr1a, respectively (Shirai et al., 2011)), within the pallium at late embryonic times (E16,5) a timepoint corresponding to the neurogenic to gliogenic switch. Analysis of the dorsal SVZ at P3 revealed profound alteration of the pattern of GFP+ cell distribution. Indeed, while overexpression of a CA Bmpr1a resulted in cells keeping a radial glia morphology and remaining in contact with the ventricular lumen, overexpression of Δ Bmpr1a led to their complete disappearance (**Fig. 6A & C**). In these mice, most GFP+ cells were located away from the ventricular surface within the SVZ and showed a round morphology, typical of progenitors. These distributions and morphologies were in agreement with a significant decrease of the number of progenitors expressing Tbr2 or Ki67 following CA Bmpr1a, while their number were consistently increased following Δ Bmpr1a overexpression (**Fig. 6B & C**). Together, these findings confirm an instructive role of TGFβ/BMP signalling on dorsal V-SVZ germinal activity through activation of the Bmpr1a receptor.

**Fig. 6.**
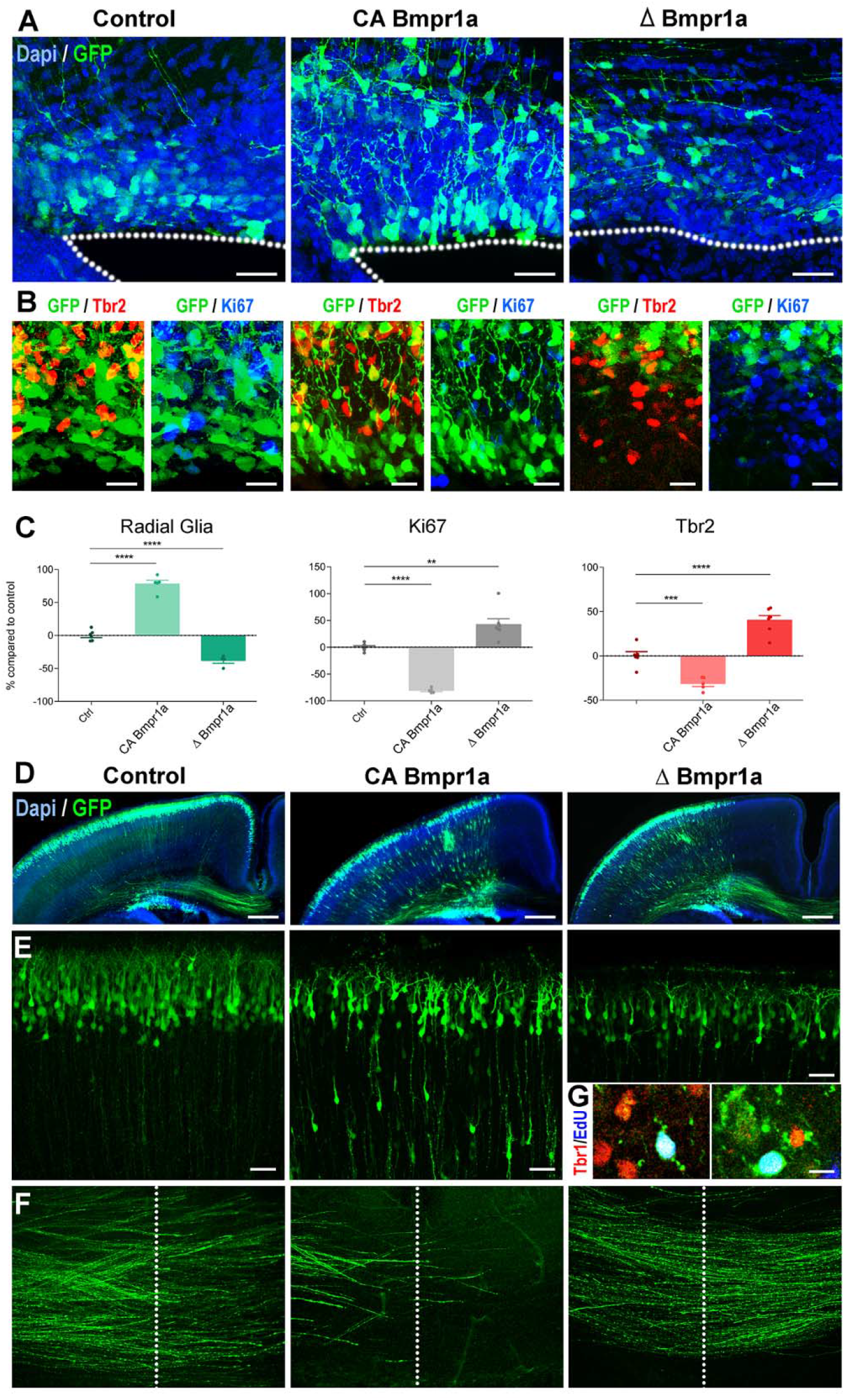
Bmpr1a manipulations modulate postnatal dorsal SVZ germinal activity. **(A)** Representative images of the dorsal SVZ at P3, electroporated with GFP (Control), constitutively active (CA) Bmpr1a and dominant negative (Δ) Bmpr1a plasmids a E16.5 The location of the ventricular lumen is indicated by a dotted line. Scale bars. **(B)** IHC for the pallial progenitor (Tbr2, red) and proliferative (Ki67, blue) markers in the 3 experimental groups. **(C)** Graphs showing the % of GFP+ electroporated cells showing RG morphology, expression of Ki67 or of Tbr2. Control values were normalized to 100% to better illustrate the consistent decrease and increase following CA Bmpr1a and Δ Bmpr1a overexpression, respectively. **(D-F)** Representative overviews of electroporated animal in the 3-experiment group (D). higher magnification of GFP positive cells within the P3 cortex (E) and of their axons within the corpus callosum (F). The midline is indicated by a dotted line. Scale bars. **(G)** Edu injection at P2 reveals the presence of Tbr1+/GFP+ neurons within the cortex following Δ Bmpr1a overexpression.

These effects were not restricted to the SVZ, but also impacted neuron migration and maturation. Indeed, overexpression of CA Bmpr1a resulted in many neurons failing to migrate to the upper most layers of the cortex (**Fig. 6D & E**). A small number of GFP+ bipolar cells were also observed within the cortical plate following Δ Bmpr1a overexpression (visible on **Fig. 6D**). EdU administration at P2 confirms that some of these cells were generated at postnatal times and expressed the neuronal marker Tbr1, suggesting an extension of the period of cortical neurogenesis in those animals (**Fig. 6G**). Finally, the maturation of upper layer neurons was also impacted, as reflected by marked differences in their axons. While these later had reached the midline by P3 in control mice, DA Bmpr1a overexpression resulted in a profound decrease of axonal growth (**Fig. 6F**). This contrasted with neurons produced following Δ Bmpr1a overexpression, which axons extended farer than in control mice, well into the contralateral hemisphere.

Thus, in agreement with an enriched expression of Bmpr1a within cells of the pallial lineage, our findings reveal that Bmpr1a signalling impact SVZ proliferation as well as the production and maturation of new-born neurons and therefore plays a key role in the closure of the period of cortical neurogenesis.

## Discussion

While the closure of glutamatergic neurogenesis was believed to mainly rely on a neurogenic-to-gliogenic switch, a recent clonal study has revealed that a large number of RGs do not undergo this switch (Gao et al., 2014). Here, we used scRNA-Seq and histological analysis of the dorsal V-SVZ to investigate at the cellular-level changes occurring in the dorsal most region at early postnatal ages. Identification and direct comparison of pallial and subpallial cell lineages within this germinal region reveal transcriptional hallmarks associated with pallial NSCs entrance into deep quiescence, their acquisition of subpallial traits, and dysregulation in their differentiation efficiency, that all converge onto a rapid closure of postnatal glutamatergic neurogenesis. Further, their comparative analysis highlight a key role for TGFβ/BMP-signalling receptor Bmpr1a in silencing pallial germinal activity after birth.

A large population of aNSCs is observed within our dataset, illustrating the sustained germinal activity observed within this region at early postnatal time points (Azim et al., 2012). We could observe an early fate priming of these cells, as illustrated by their clear segregation in cell clusters expressing TFs known to instruct neurogenesis, astrogenesis and oligodendrogenesis. Further, our results support the co-existence of both unipotent and bipotent NSCs, as reflected by the co-expression of neurogenic and oligodendrogenic TFs (e.g. Eomes and Olig2, not shown) in some aNSCs and TAPs. These observations are in line with the early priming of adult NSCs (Llorens-Bobadilla et al., 2015), as well as with recent clonal fate-mapping studies within the postnatal V-SVZ (Figueres-Oñate et al., 2019). Thus, while at the population level, NSCs appear to act in a rather homogeneous manner, analysis at the single-cell level reveals a level of unsuspected heterogeneity.

Remarkably, RNA velocity reveals several trajectories emerging from aNSCs, the most noticeable driving them toward quiescence. Thus, although major transcriptional differences between the dorsal and lateral V-SVZ have been shown by previous studies (Azim et al., 2015, 5), our work highlights novel important differences in NSCs dynamics, in particular in the timing of induction of their quiescence. Indeed, while original studies have pointed out an embryonic origin of adult NSCs (i.e. “set-aside” model, (Fuentealba et al., 2015; Furutachi et al., 2015), our results indicate that this rule does not apply to the dorsal V-SVZ. Indeed, both our transcriptomic and histologic analyses reveal an entrance into quiescence at perinatal age. These conclusions are in line with fate mapping of embryonically born qNSCs in the H2B-GFP mouse following E9.5 induction, which revealed that GFP retaining cells are observed within the lateral, but not the dorsal most regions of the juvenile V-SVZ (Furutachi et al., 2015). Similarly, our results confirm that BrdU injection at E14.5 results in the detection of numerous positive cells within the lateral and ventral V-SVZ (data not shown), while no cells are observed within the dorsal V-SVZ (Fuentealba et al., 2015). Thus, while a population of qNSCs appears to diverge from embryonically active ones within the ganglionic eminence and to persist at postnatal ages in the lateral and ventral V-SVZ, those from the dorsal V-SVZ enter quiescence around birth, in line with a “continuous model” recently proposed within the hippocampus (Berg et al., 2019). Knowing the importance of the tight regulation of qNSCs/aNSCs balance in maintaining germinal activity, it is tempting to speculate that this temporal difference results in distinct states of germinal activity sustainability within the SVZ. Thus, while NSCs derived from the subpallium remain germinally active in producing various olfactory bulb interneurons throughout life, NSCs derived from the pallium declines rapidly in their capacity to produce glutamatergic periglomerular cells (Brill et al., 2009; Winpenny et al., 2011), then CR+ interneurons (Kohwi et al., 2007), possibly through their rapid exhaustion or induction of irreversible quiescence.

NSCs expressing pallial or subpallial markers coexist within the postnatal V-SVZ dorsal domain, allowing for the first time, their direct comparison. Interestingly, distinct levels of quiescence characterize these two cell populations. Indeed, parallel comparison of pallial and subpallial quiescence trajectories, reveals the synchronized and persistent expression in subpallial qNSCs of genes involved in biological functions known to be important for germinal activity maintenance. For instance, genes involved in extracellular matrix (ECM) reorganisation and regulation of cell adhesion, are observed within subpallial qNSCs, in agreement with the importance of NSCs interactions with their microenvironment in regulating quiescence/activation balance (Codega et al., 2014; Kazanis et al., 2010). This is illustrated by the marked enrichment of Vcam1 which was shown to act in adult NSCs induction and maintenance (Hu et al., 2017; Kokovay et al., 2012).

Although also observed at the early stages of the pallial qNSCs trajectory, these gene sets appear to gradually vanish from pallial NSCs, suggesting their gradual loss of anchorage within the ECM and progressive isolation from both the vasculature and CSF. This is supported by our observation that most pallial qNSCs (Hopx-derived Sox2+/BrdU-, see below) can be seen at P21 at some distance from the ventricular lumen as well as in rostral SVZ regions, where the ventricle has collapsed. Deeper quiescence is further supported by multiple observations, including a reduced level of P27 (Cdkn1b) and Egfr expression, which defines acquisition of a primed GO state (Marqués-Torrejón et al., 2021) and entrance in proliferation (Pastrana et al., 2009), respectively.

Our results highlight Hopx as a marker of pallial qNSCs. Interestingly, Hopx was identified as a quiescent stem cell marker in intestines (Takeda et al., 2011) and hematopoietic stem cells (Lin et al., 2020) in mice, as well as in dentate gyrus NSCs (Berg et al., 2019), another region of postnatal glutamatergic neurogenesis. The restricted Hopx expression within pallial NSCs allowed us to fate map those cells at early postnatal stages (P1). Interestingly, at this early stage, a large proportion of Hopx+ cells are non-proliferative as revealed by the absence of BrdU incorporation. Further, at least a portion of those cells persist within the dorsal V-SVZ for at least 3 weeks after birth (our observations) and probably later (see for example (Cebrian Silla et al., 2021)). Interestingly, Hopx transcripts remain detectable within adult qNSCs where they show a >33-fold enrichment when compared to aNSCs (Codega et al., 2014). Fate mapping of Hopx-expressing cells at 2 months results in no neurons being labelled in the OB (Li et al., 2015), further supporting deep, irreversible quiescence. It is unclear if Hopx contributes directly to the induction of deep quiescence. Indeed, a comparative GSEA analysis of Hopxhigh and Hopxlow qNSCs fails to reveal significant gene set induction in Hopxhigh cells, while its low expression correlates with priming for activation, as reflected by induction of ribosomal and mitochondrial biogenesis transcripts. Thus, while Hopx knock-out disrupts stemness and quiescence of hematopoietic stem cells in mice (Lin et al., 2020), its role is likely to be more complex within the postnatal V-SVZ. Indeed, previous experiments aimed at investigating the consequences of Hopx GoF and LoF within the postnatal V-SVZ, failed to demonstrate a marked effect on germinal activity, although induction of quiescence was not investigated (Zweifel et al., 2018).

The molecular mechanisms underlying Hopx+ NSCs quiescence appear to be at least partially related to regulation of TGFβ versus Wnt-signalling. Hopx has been shown to modulate primitive hematopoiesis by inhibition of Wnt-signalling (Palpant et al., 2017), and modulation of TGFβ/Wnt-signalling by Hopx has been shown to occur in the developing heart (Jain et al., 2015), as well as during endothelial development and primary hematopoiesis (Palpant et al., 2017). Interestingly, our transcriptional results suggest a persistent activation of TGFβ-signalling in pallial qNSCs. Further, the transcriptional profile of pallial qNSC resembles those of NSCs exposed to BMP4 in vitro, which in the absence of FGF2 result in induction of deep quiescence (Marqués-Torrejón et al., 2021; Mira et al., 2010). BMPs belong to the superfamily of TGFβ cytokines and exert a plethora of effects in the nervous system, ranging from dorsoventral patterning to induction of the neurogenesis to gliogenesis switch (Chen and Panchision, 2007). Thus, TGFβ-signalling may play a pleiotropic role in pallial lineage germinal activity closure, by acting in both astrogenesis in concert with JAK-STAT signalling (Nakashima, 1999) and induction of deep quiescence for cells of the pallial lineage. Our observation that stemness transcripts are retained, while GFAP expression remains absent at P21 in the Hopx progeny within the SVZ, suggests BMP signals do not drive terminal astrocyte differentiation, but rather impose quiescence in at least a fraction of pallial NSCs (Marqués-Torrejón et al., 2021). This is in line with a role of TGFβ-signalling in regulating the temporal identity and potency of NSCs in other regions of the CNS, such as the hindbrain (Dias et al., 2014) and the retina (Kim, 2005). Identification of TGFβ family members and receptors within the dorsal SVZ niche, suggests that BMP4 and 5 are involved, which are secreted by OPCs/newly-formed OLs and NSCs themselves, respectively (Azim et al., 2017). Furthermore, an enrichment for Bmpr1a expression is observed in pallial qNSCs. Bmpr1a is responsible for triggering the canonical BMP pathway by phosphorylating the SMAD1 protein, which translocation is associated with the upregulation of target genes, such as Id1-4 (Mira et al., 2010), all of which show higher expression in pallial NSCs and are known to block the action of pro-differentiation bHLH transcription factors (Blomfield et al., 2019). This elevated expression of several inhibitors of transcription, correlates with a mild reduction in their transcriptional activity, possibly by the recruitment of class I histone deacetylases (Hdacs) as observed in other tissues (Trivedi et al., 2010). This is in line with our gain and loss of function experiments, which demonstrate that manipulating Bmpr1a results in profound changes in the germinal activity of the dorsal SVZ. While activation of Bmpr1a results in a persistence of radial glial cells at postnatal times, its inactivation has the opposite effect by releasing proliferating and glutamatergic (i.e. Tbr2+) progenitors’ production.

Although the entrance of pallial NSCs in a state of deep quiescence may solely explain the silencing of pallial germinal activity early after birth, other mechanisms appear to participate in the rapid decline of postnatal glutamatergic neurogenesis. For instance, it was recently proposed that some pallial progenitors acquire subpallial traits at postnatal timepoints (de Chevigny et al., 2012; Zhang et al., 2020), which correlates with the demonstration of a subpopulation of olfactory bulb gabaergic neurons deriving from Emx1 progenitors at postnatal ages (Kohwi et al., 2007) under the influence of Shh signalling which activity rises in the dorsal V-SVZ at postnatal stages (Tong et al., 2015). These findings are in line with our observation of a significant number of dual cells, co-expressing markers of both the pallial and subpallial lineages, within our dataset, as well as by our fate mapping of E14.5 pallial NSCs. Our results however show that this “reprogramming” of pallial NSCs is however largely incomplete, with a subpopulation of TAPs and nascent neurons presenting a clear glutamatergic identity observable by scRNA-Seq until at least P22, in line with our previous histological and fate mapping studies (Azim et al., 2012, 2014; Donega et al., 2018). Their transcriptional comparison with surrounding gabaergic TAPs reveal a reduced proliferative capacity, as well as a failure of expression of genes necessary for tangential migration toward the OB. Further, their transcriptional analysis at distinct postnatal time (i.e. P2 to P22) support their rapid failure to undergo efficient differentiation in line with their persistent expression of Id proteins and Bmpr1a. These observations are in line with observations made following Bmpr1a gain and loss of function. Thus, while activation of Bmpr1a at late embryonic timepoints results in many neurons failing to migrate and mature, its inactivation results in a more rapid development of interhemispheric axonal projection, as well as in a prolongation of the period of cortical neurogenesis as supported by the observation of EdU+/Tbr1+ neurons produced at postnatal stages.

Altogether, our results represent the first characterization of the postnatal dorsal SVZ by scRNA-Seq. By allowing a direct comparison of pallial and subpallial cell lineages, our work shed new light on the events that ultimately result in the rapid closure of the period of glutamategic neurogenesis while gabaergic neurogenesis persists throughout life. In particular, the enrichment of Bmpr1a expression within cells of the pallial lineage appears to play a key role in synchronizing quiescence induction and silencing of neuronal differentiation to rapidly silence pallial germinal activity after birth.

## Materials and Methods

### Animals and Ethics

Mice used in this study were OF1 wild type (Charles Rivers, France) and HopxCreERT2 (Takeda et al., 2011). In HopxCreERT2 animals, subcutaneous tamoxifen (Tam; SIGMA) administration (1 mg per pup) was performed at P1 (i.e. 24 hours following birth). All animal experiments were performed in accordance with European requirements 2010/63/UE had have been approved by the Animal Care and Use Committee CELYNE (APAFIS #187 & 188). Mice were group-housed, with food and water ad libitum, under 12 h light–dark cycle conditions.

### Tissue preparation for single-cell RNA sequencing

#### Single-cell isolation

The dorsal subventricular zone (dV-SVZ) from P12 mice of 2 independent experiments, as well as of P2 and P22 mice, were dissected. During all stages of the dissociation protocol, the tissue was kept in artificial CSF solution containing 125 mM NaCl, 2.5 mM KCl, 1.25 mM NaH2PO4, 26 mM NaHCO3, 17 mM glucose, 1.25 mM CaCl2 and 1 MgCl2. The small dissected tissues were incubated in Papain during 20 minutes at 37°C. Following enzymatic digestion, cells were centrifuged for 5 minutes and the pellet was resuspended with DNAase/ovomucoid inhibitor according to manufacturer’s protocol (Worthington). Cells were then centrifuged and resuspended on ice in Leibovit’z L-15 medium supplemented with 0.1% of bovine serum albumin. The cell suspension was finally filtered through a 70-μm cell strainer to remove aggregated cells.

#### FACS

Viable cells (DAPI-and Hi Draq5+) were then sorted using BD FACS Aria sorter. 30 000 cells were collected per replicate in ice-cold PBS.

### Single-cell RNA sequencing

Cell suspension (300 cells per µl) was added to 10x Genomics Chromium Single-Cell Controller (10x Genomics) to achieve 7000 encapsulated cells per replicate. The next steps for cDNA synthesis and library preparation were done following the manufacturer’s instructions (chemistry V3). Libraries have been sequenced independently using the Novaseq 6000 platform (Illumina) in order to target 100k reads per cell. Cell Ranger version 3.0.1 was used to align reads on the mouse reference genome GRCm38 mm10 and to produce the count matrix.

### Analysis of single-cell RNA-sequencing data

#### Quality control and filtering

We filtered 11 298 cells (P12 dataset) 4037 cells (P2 dataset) and 4254 cells (P22 dataset), based on two-quality control criteria: the number of genes per cell and the fraction of counts from mitochondrial genes per cell. Cells with <2500 genes or >7500 and with >10% mitochondrial genes fraction were removed.

#### Clustering analysis

We first focused our analysis on the P12 replicates. Filtering and Data analysis were performed using the R package Seurat (version 3.1) (Stuart et al., 2019). First, genes expressed in less than 3 cells were removed in each dataset. Gene expression was normalized using the standard Seurat workflow and the 2000 most variable genes were identified. Then, the 2 batches were integrated using the integration function of Seurat and anchor genes were scaled without any type of regression and used for PCA at 50 dimensions. We then performed preliminary clustering with permissive parameters to identify and remove low-quality clusters. We only removed one cluster of 19 cells that expressed fewer genes compared to average. The remaining cells were subjected to second-level clustering (20 PCs; resolution = 0.5) yielding 15 final clusters (full_identity), merged into 9 clusters (simplified_identity). The clusters were visualized in two dimensions using the RunUMAP() function (minimum distance = 0.3; n_neighbors = 30 L; UMAP.method = ‘UMAP-learn’; metric = ‘correlation’). We then performed differential expression analysis and categorized each cluster as one of the following major cell types. Identification of cell clusters within the P2 and P22 datasets was achieved based on P12 cell types using MapQuery() function. For comparative/quantitative analysis of P2, P12 and P22 datasets (e.g. **Fig. 2O**), NBs and OLs were excluded to avoid possible influences from the microdissection that may result in the inclusion of some RMS or overlying corpus callosum.

### Sub clustering

For sub clustering of aNSCs, corresponding to 1514 cells, a resolution of 0.5 was used leading to the appearance of 4 subclusters. For sub clustering of aNSC3, representing 569 cells, a resolution of 2 was used. Finally, for sub clustering of the neuronal trajectories, corresponding to 1029 cells, a resolution of 0.75 was used.

### Transcriptional criteria for cell type identification

We defined the various cell types based on marker combinations (normalized Logged counts): qNSCs (Slc1a3>1 & Prom1 >0.6 & Egfr<0.1 & Foxj1<0.1); aNSCs (Egfr>1 & Ascl1>1 & Dlx1<0.01 & Dlx2<0.01); TAPs (Egfr>1 & Ascl1>1 & Dlx1>0.01 & Dlx2>0.01 & Sp8<0.5 & Gad1<0.5 & Gad2<0.5); NBs (Dcx>1 & Cd24a>0.5 & Nrxn3>0.5); OLs (Pdgfra>0.5 & So×10>0.5) and Astrocytes (S100b>0.5 & Aqp4>2). Pallial cells were defined as follow: (Emx1>0.25|Neurod1>0.25|Neurod6>0.25|Neurog2>0.25|Tbr1>0.25)&(Gsx2<0.25&Dlx6<0.25& Dlx5<0.25&Sp9<0.25&Dlx2<0.25&Gad1<0.25) and Subpallial cells were defined as follow: (Gsx2>0.25|Dlx6>0.25|Dlx5>0.25|Sp9>0.25|Dlx2>0.25|Gad1>0.25)&Emx1<0.25&Neurod1<0.2 5&Neurod6<0.25&Neurog2<0.25&Tbr1<0.25).

### RNA Velocity

We predicted the direction of transcriptional changes using the RNA velocity framework, which estimates the gene expression dynamics from exonic and intronic expression. We first annotated spliced, unspliced and ambiguous reads using the run10x command (velocyto.py). RNA velocities were then predicted using the R package scVelo (La Manno et al., 2018) based on the 2000 most variable genes and 30 PCs and 30 neighbours. Estimated RNA velocities are represented on the UMAP with streamlines (scv.pl.velocity_embedding_stream). The direction of the arrows indicates the estimated future state of the current cells. Long arrows correspond to large gene expression changes. We determine RNA velocity using stochastic and dynamical models.

### Data visualization

To display the gene expression, the preprocessed UMI matrix was normalized with the function library.size.normalization of the R package Magic (van Dijk et al., 2018). The dropout corrected data were displayed on the Seurat FeaturePlots with the viridis scale colors only. Otherwise, the grey-red scaled FeaturePlots illustrate the original non-corrected UMI matrix.

### Pseudotime

Both pallial and subpallial quiescent cell lineages have been isolated and the R package slingshot used to calculate the pseudotime values using the cluster aNSC3 as root of the trajectory (**Fig.3E**).

### Differential expression analysis

To identify enriched genes, we performed a non-parametric Wilcoxon rank-sum test using the FindMarkers() function from Seurat. p-value adjustment is performed using bonferroni correction based on the total number of genes in the datasets. Genes with a p-value <0.05, at least 0.25 average fold change (log scale) and at least 25% cluster-specific detection (percentage of cells expressing a particular gene in a cluster) were defined as enriched genes for each cluster.

### Cell cycle assignment

For each cell, we computed a score based on its expression of G2/M and S phase markers; cells that express neither of these markers are likely not cycling or are in G0/G1 phase. The scoring strategy is described in (Kowalczyk et al., 2015). We used Seurat’s implementation of this strategy in the CellCycleScoring() function.

### Identification of differentially expressed genes

Identification of differentially expressed genes was based on the following criteria min.pct = 0.1, logfc.threshold=0.25.

### Identification of transcription factors and transcriptional regulators

Differentially expressed genes were compared to gene sets “DNA Binding” (GO:0003677) et “Transcriptional activity” (GO:0140110).

### Gene set enrichment

Over-representation analyses were performed using the R package “Cluster Profiler” (https://guangchuangyu.github.io/2016/01/go-analysis-using-clusterprofiler/) (Yu et al., 2012). DEG were extracted by using the following criteria (Logfc.threshold=0.25, min.pct=0.1). GSEA analyses (Subramanian et al., 2005) were performed by using the GSEA software v4.1.0 [Built 27] from the board institute (GSEA (gsea-msigdb.org)), using the following curated gene sets databases: Hallmarks “hall.v7.3.symbols.gmt”, Kegg “c2.cp.kegg.v7.3.symbols.gmt”, Reactome “c2.cp.reactome.v7.3.symbols.gmt”, Gene Ontology c5.go.bp.v7.3.symbols.gmt”. Obtained results were exported in cytoscape 3.8.1 for visualization and analysis. (www.cytoscape.org).

### ClusterMap analysis

A correlative analysis was performed using the R package “ClusterMap”. It provides a way for automatically and unbiasedly matching sub-groups based on a list of marker genes detected in the analysis. ClusterMap can generate circular graphs, called circos plots, between several datasets. It identifies the similarity of clusters, based on transcriptional similarities, and represents them as an arc linking one or more clusters together. First, the gene list for each sub-cluster in each sample is used to match the sub-cluster between samples. The file containing the gene list can be the direct output of the Seurat package’s “FindAllMarkers” function. The arcs in the circos plot indicate the subgroups’ relationships. The width of the gray or pink sectors represents the proportion of cells in each sample. Lastly, the degree of similarity between the matched groups is shown by the transparency of the hue of the arcs. Similarity is a measure of the percentage of overlapping genes between groups (Gao et al., 2019).

### Integration of multiple datasets

Integration was done by identification of cells that are in the same biological state (anchors) using the function FindIntegrationAnchors() of the Seurat R package. The function IntegrateData() was then applied with the previous anchor set to perform dataset integration using the rpca (Reciprocal PCA) approach. The following datasets were used for integration: GSE123335 dataset corresponding to embryonic (E14.5) and neonatal (P0) pallium and overlying cortex (Loo et al., 2019); GSE134918 dataset corresponding to P56 and P70 V-SVZ and olfactory bulb (Mizrak et al., 2020).

### Label retaining protocol

A single injection of BrdU (50mg/kg, i.p.) was made in pregnant mice at E14.5 or in newborn mice (P1, P3, P5). Mice were sacrificed at P21.

### Histology and Immunostainings

Mice were sacrificed by an intraperitoneal overdose of pentobarbital followed by perfusion with Ringer’s Lactate solution and 4% paraformaldehyde (PFA) dissolved in 0.1 M phosphate buffer (PB; pH 7.4). Brains were removed and postfixed for 12–48 hr at 4°C in 4% PFA and sectioned in 50-µm thick coronal serial sections. When necessary, antigen retrieval was performed for 20 min in citrate buffer (pH 6) at 80°C, then cooled for 20 min at room temperature and washed in 0.1 M PB. Immunostaining was performed as previously described (Donega et al., 2018; Zweifel et al., 2018).

Blocking was done in TNB buffer (0.1 M PB; 0.05% Casein; 0.25% Bovine Serum Albumin; 0.25% TopBlock) with 0.4% triton-X (TNB-Tx). Sections were incubated overnight at 4°C with gentle shaking the following primary antibodies in TNB-Tx. The following primary antibodies were used for immunohistochemical procedures: Rabbit anti-Hopx (1:400; Santa Cruz; sc-30216); Mouse anti-Hopx (1:400; Santa Cruz; sc-398703); Goat anti-DCX (1:500; Santa Cruz; sc-8066); Mouse anti-GFAP (1:500; Millipore; MAB3402); Goat anti-Mcm2 (1:300; Santa Cruz; sc-9839); Mouse anti-Sox2 (1:100; Santa Cruz; sc-365823); Guinea Pig anti-S100b (1: 2000; synaptic system); Rat anti-Tbr2 (1:1000; Invitrogen; 14-48-75-82); Rabbit anti-Tbr2 (1:1500; Millipore; AB2283) Rabbit anti-GSX2 (1:2000; Millipore; ABN162); Rabbit anti-S100b (1:5000; Swant); Rat anti-BrdU (1:1000; Abcam; ab6326); Mouse anti-BrdU (G3G4) (1:1000; DSHB; 7/5118), Rabbit anti-Tbr1 (1:500; Abcam; AB31940). Following extensive washing in 0.1 M PB with 0.4% triton-X (PB-Tx), sections were incubated with appropriate secondary antibodies conjugated with Alexafluor 488, 555 or 647 (1:500; Life Technologies) for 2 hrs at room temperature. Sections were washed and counterstained with Dapi (1:5000; Life Technologies; D1306).

### Electroporations

The following plasmids were used in this study: pCX-GFP (kind gift of X Morin, ENS, Paris; France); pPB-CAG-EmGFP (VB161220-1119syh; VectorBuilder Inc., Cyagen Bioscience, Santa Clara, California, USA); pCMV-hyPBase (kind gift of Laura LLpez-Mascaraque; Instituto Cajal, Madrid, Spain), pcDNA3-CAG-dnALK3 and pcDNA3-CAG-caALK3 (Kind gift of Kohei Miyazono, University of Tokyo, Japan, corresponding to dominant-negative and constitutively active forms of Bmpr1a, respectively). Plasmids were purified using the EndoFree Plasmid Kit according to the manufacturer’s protocol (Qiagen; 12362). Plasmids were re-suspended to a final concentration of 5 µg/µl. For postnatal fate mapping of pallial radial glial cells, co-electroporation of pPB-CAG-EmGFP and pCMV-hyPBase plasmids was done at E15.5. For Bmpr1a manipulation, electroporation was directed towards the embryonic pallium at late embryogenesis time, i.e. E16.5, as previously described (Meyer-Dilhet and Courchet, 2020). EdU (50mg/kg) was injected 24hrs before sacrifice (i.e. P2).

## Figures

Venn diagrams. Venn diagrams were done by using “Multiple List Comparator” available online (https://molbiotools.com/listcompare.php).

## Acknowledgments

We are grateful to Kasum Azim for his critical comments onto the manuscript, Timothy Capeliez for help with the histology, as well as Julien Courchet and Geraldine Meyer-Dilhet for their help with the electroporation experiments. We thank the Single-Cell Bioinformatics Platform of the Labex Cortex, the Centre de Recherche en Cancérologie de Lyon (CRCL), as well as the Centre Leon Berard (CLB) in Lyon, France, for their help in producing the transcriptional datasets presented here, as well as for their guidance in performing the analysis.

## Author contributions

Conceptualization: GM, LF, OR

Methodology: GM, LF, EB, ET, SZ, CP, DJ, HHV, CH, OR

Investigation: GM, LF, EB, ET, OR

Supervision: OR, GM

Writing—original draft: OR

Writing—review & editing: OR, GM, LF, HHV, CH, CP, DJ

## Data and materials availability

The accession numbers for the data reported in this paper are GEO: GSE XXXXXX.

## Supplementary Materials

Please see the Supplementary Materials document.

**Fig. S1.**
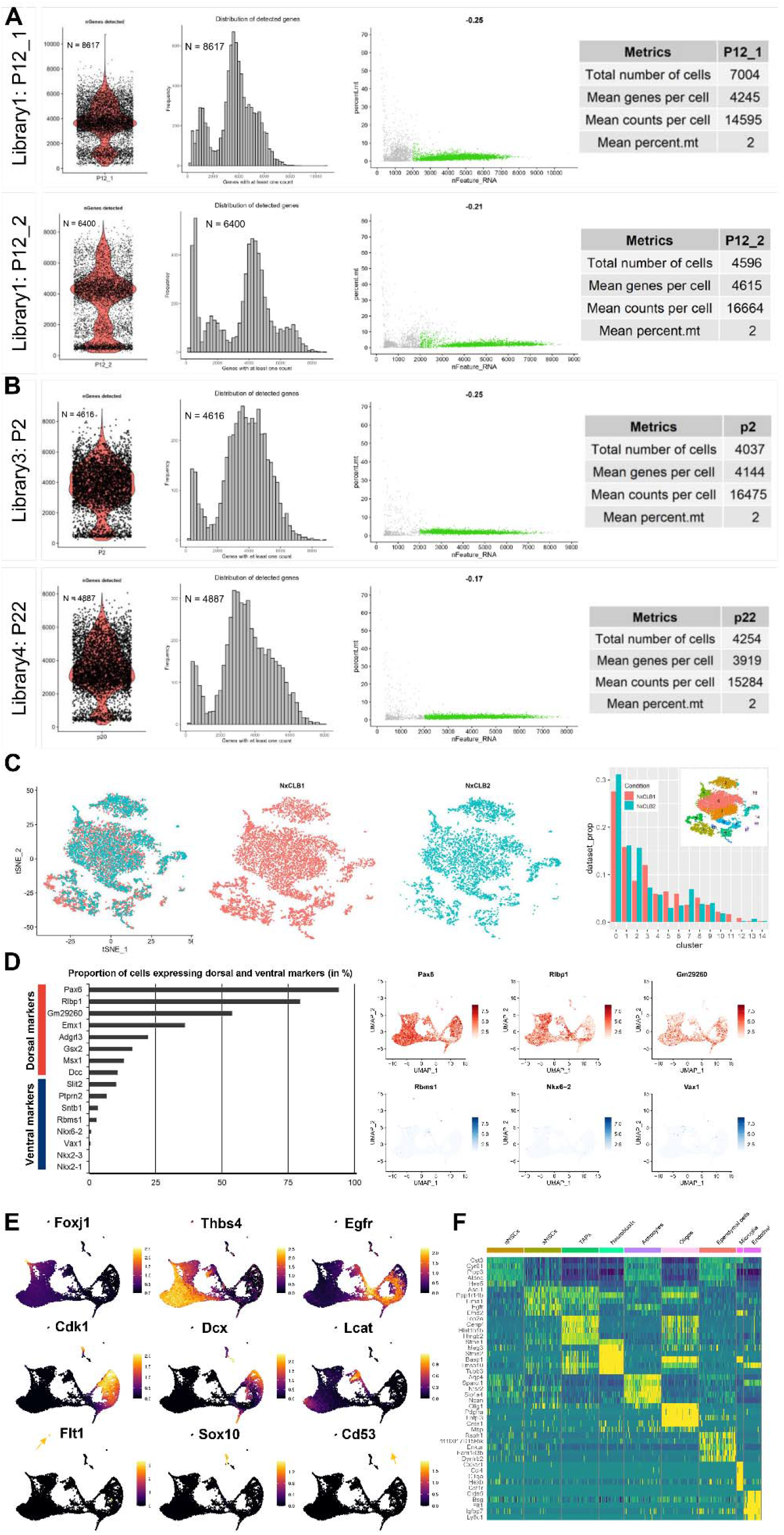
Data Metrics and Markers, *related to Fig 1*. (A) Summary of Metrics of the P12 replicates. Cells to be analyzed are selected based on the following criteria: percent.mt<10% and number of genes expressed >2000 & <8000. **(B)** Summary of Metrics of the P2 and P22 additional datasets. Cells to be analyzed are selected based on the following criteria: percent.mt<10% and number of genes expressed >2000 & <8000. **(C)** The dimensionality reduction technique tSNE shows the great overlap of the P12 replicates. Their homogenous contributions to the 15 clusters are shown in percentage. **(D)** Percentage of cells expressing dorsal and ventral SVZ markers, and feature plots of most selected genes illustrating the precision of the microdissection approach. **(E)** Markers of the main cell types observed in the dataset are shown as FeaturePlots. **(F)** Heatmap depicting expression of top 5 markers distinguishing cell types.

**Fig. S2.**
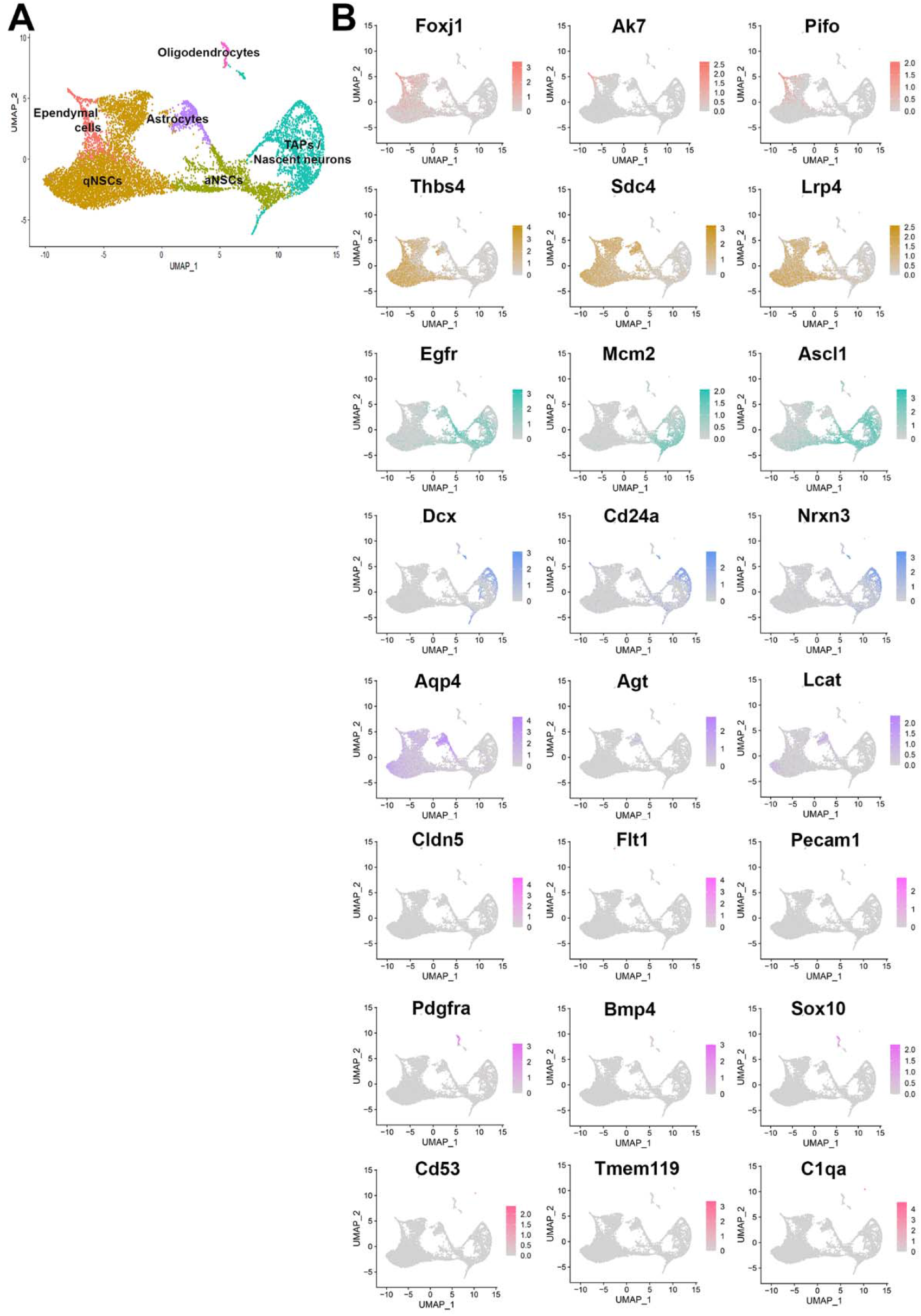
A core set of “generic genes” defines distinct cellular states, *related to Fig 1*. **(A)** UMAP with simplified identity annotation. **(B)** Feature plots of markers used to define cell types. Note the overlap of aNSCs markers with TAPs.

**Fig. S3.**
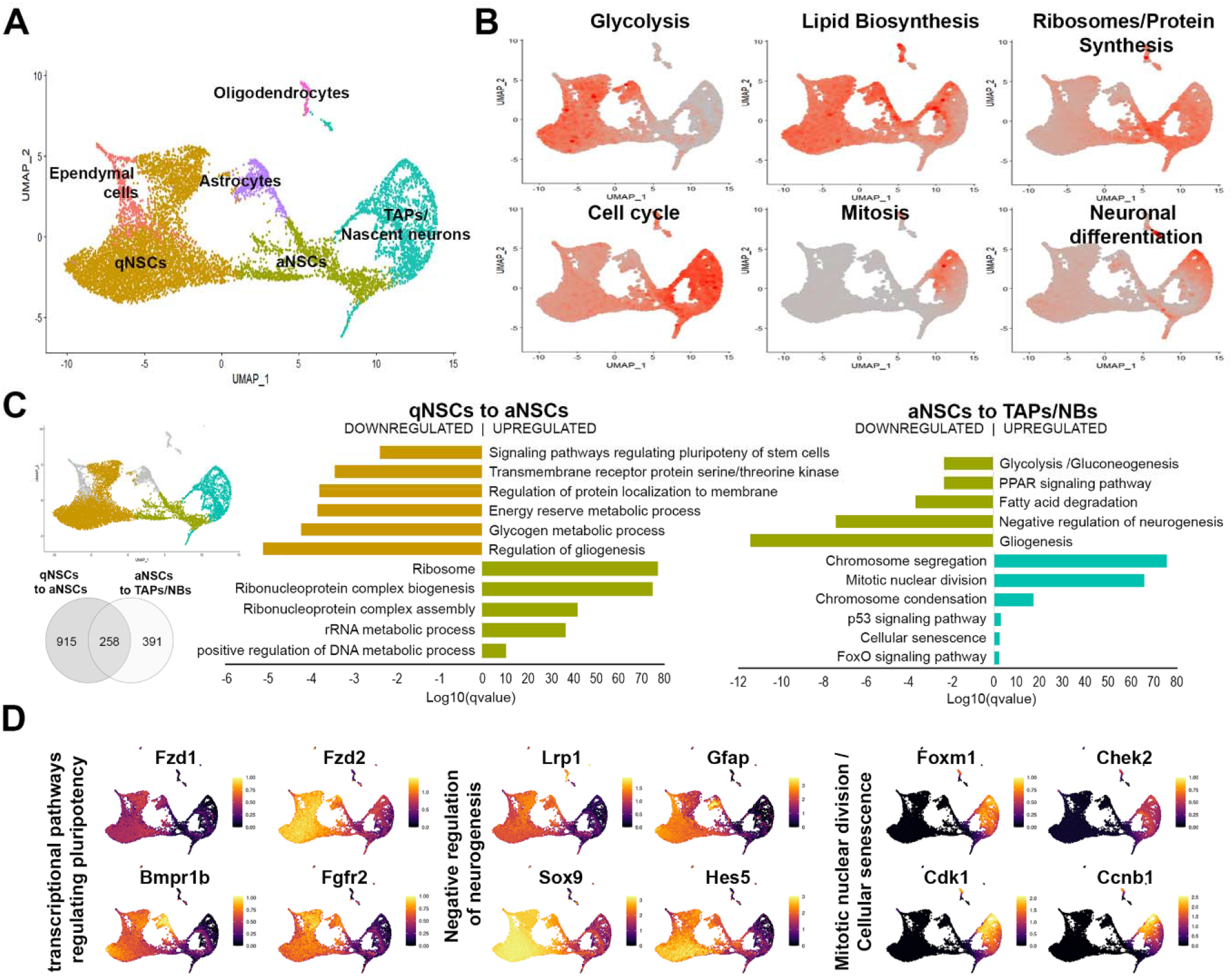
A core set of “generic genes” defines distinct cellular states, *related to Fig 1*. **(A)** UMAP with simplified identity annotation. **(B)** Percentage of expression of top 20 genes associated with previously known markers reflecting state transitions along the neurogenic lineage (Llorens-Bobadilla et al., Cell Stem Cell, 2015). Dormancy is associated with high glycolytic and lipid metabolism. Lipogenesis has recently emerged as a key metabolic pathway in hippocampal NSC maintenance (Knobloch et al., 2013). qNSCs share many markers (i.e., Aldh1l1 and Gjb6) and molecular features (i.e., high glycolytic activity). **(C)** Gene ontology and pathway analyses on “generic genes” allowing transition between qNSCs, aNSCs and their progeny (TAPs and neuroblasts). **(D)** Select genes from regulated gene sets illustrating the dynamics of gene expression during differentiation progression. Note the persistent downregulation of genes involved in gliogenesis as well as Glycolytic metabolism to allow differentiation progression.

**Fig. S4.**
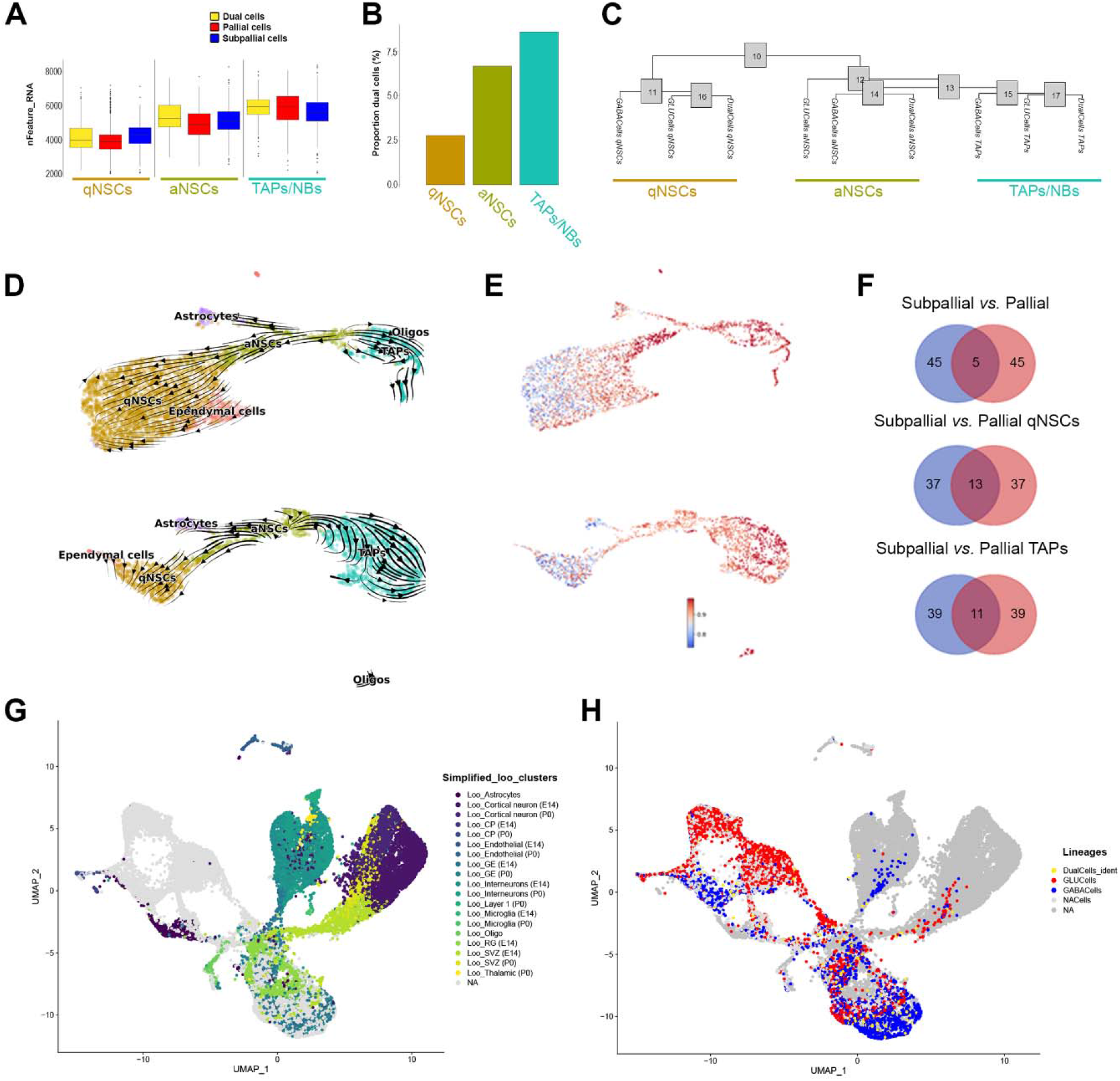
Pallial and subpallial lineages coexist within the dorsal SVZ, *related to Fig 2 & 3*. **(A-C)** Dual cells are not doublets as confirmed by a similar average number of detected genes compared to pallial cells and subpallial cells (A). Dual cells proportions within SVZ cell types (B). Hierarchical tree shows gradual identity switch of dual cells from a pallial to a subpallial identity (C). **(D-E)** Pseudotime calculated by RNA velocity (stochastic model – scVelo) highlights the trajectories cells of the pallial (top) and subpallial (bottom) lineages (D), with high confidence values (E). **(F)** Ven diagram illustrating the minimal overlap of top 50 genes contributing to the velocity calculated in both lineages, qNSCs or TAPs. **(G-H)** UMAP plots of integrated datasets presented in Fig 3A, with identity of Loo datasets (G), and of pallial and subpallial cells from current P12 dataset (H).

**Fig. S5.**
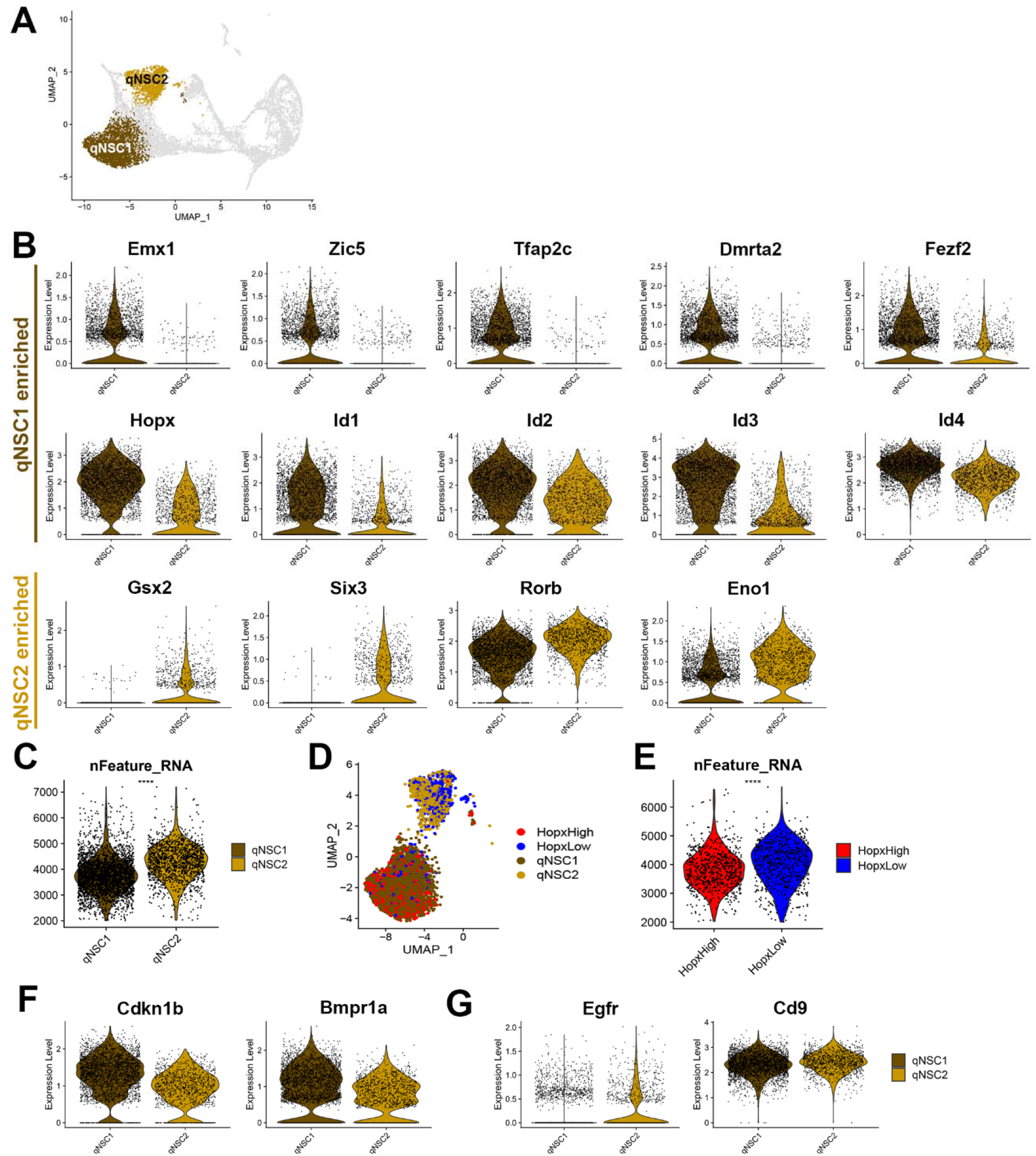
Pallial and subpallial qNSCs differ in their transcriptional profile and transcript content, *related to Figure 4*. **(A)** UMAP highlighting qNSC1 and qNSC2 subclusters. **(B)** Violin plots illustrating expression of transcription factors/transcriptional regulators enriched in qNSC1 and qNSC2. **(C)** Violin plots illustrating higher transcript content of qNSC1 when compared qNSC2. **(D)** UMAP plot showing enrichment of Hopx^Hight^ cells in qNSC1, while Hopx^Low^ cells are mainly associated to qNSC2. **(E)** Violin plot showing accordingly higher transcript content in Hopx^low^ cells when compared to Hopx^high^ cells. **(F-G)** Violin plots illustrating the enrichment of genes associated to deep quiescence (F) or primed quiescence (G), as defined by Marqués-torrejón, M. Á, Nat Comm 2021.

**Fig. S6.**
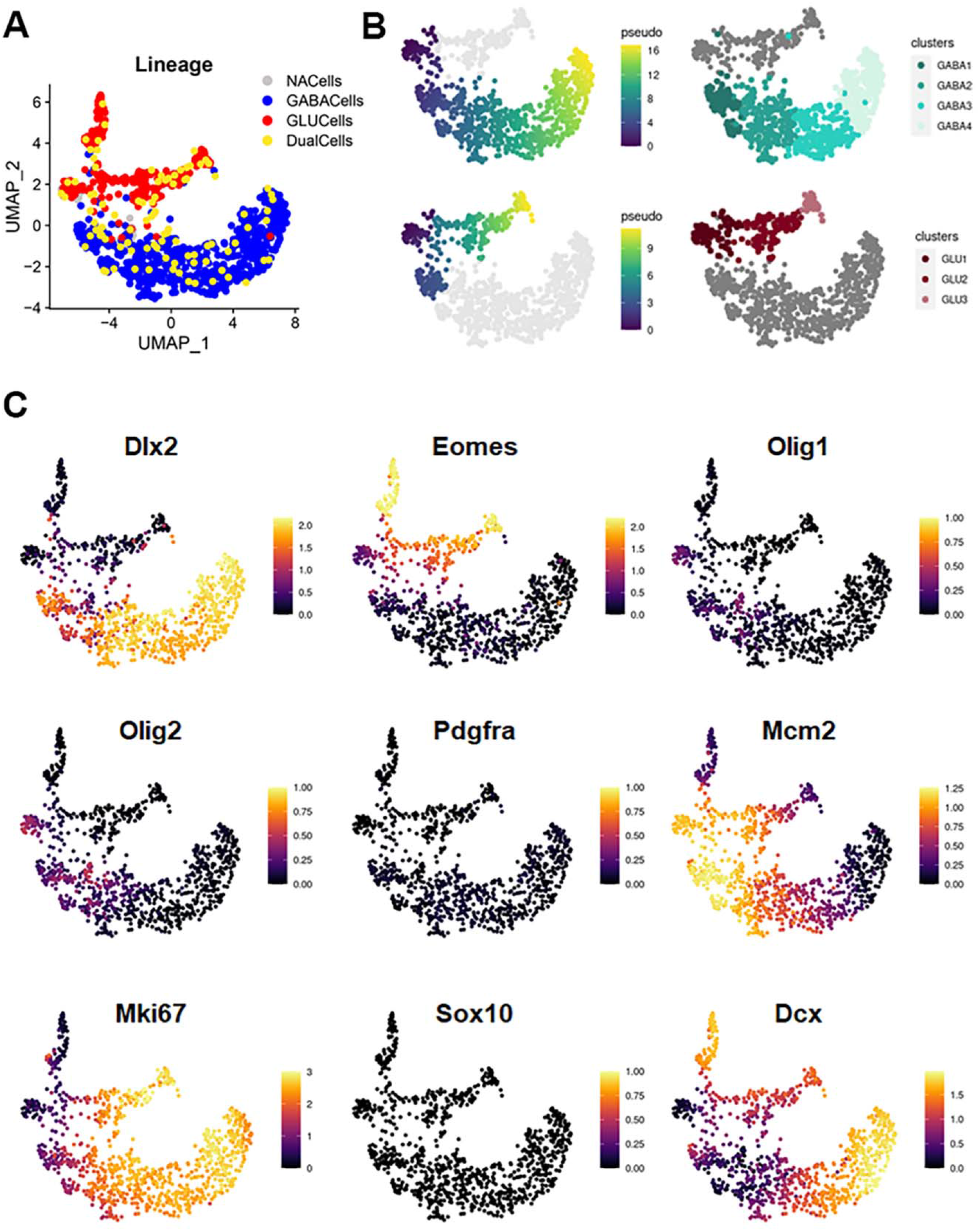
characterization of GLU and GABA cells within the neuronal trajectory, *related to Figure 5*. **(A)** UMAP highlighting distribution of GLU, GABA and Dual cells within the neuronal trajectory. **(B)** Pseudotime analysis within cycling GLU and GABA cells, and related subclusters. **(C)** Violin plots illustrating expression of various cell types or cell status markers.

**Data S1:** <u>ORA_subclusters_aNSCs.</u> Table of gene ontology analysis (simplified biological processes) for genes enriched in all aNSCs subclusters. Related to figure 2C.

**Data S2:** <u>aNSC3 subcluster markers.</u> Table of genes enriched in aNSC3 subclusters. Related to figure 2D-E.

**Data S3:** <u>GSEA_pallial vs subpallial qNSC trajectories.</u> Table of curated gene set enrichment analysis performed at 1^st^ and 2^nd^ transition steps of the pallial and subpallial qNSCs trajectories. Related to figure 3E-F.

**Data S4:** <u>ORA_Hopxhighandlow qNSCs:</u> Table of overrepresentation analysis for GO simplified biological processes, KEGG and Reactome pathways in qNSCs showing high or low levels of Hopx expression. Related to figure 4H.

**Data S5:** GSEA_Hopx^high^ vs low_qNSCs. Table of ranked gene list and GSEA reports for gene sets enriched in Hopx^high^ and Hopx^low^ qNSCs. Related to figure 4I.

**Data S6:** <u>ORA_GLU vs GABA cells:</u> Table of gene ontology analysis (simplified biological processes) for genes enriched in GLU or GABA cells. Related to figure 5G-H.

